# Cellular and behavioral effects of altered Na_V_1.2 sodium channel ion permeability in *Scn2a^K1422E^* mice

**DOI:** 10.1101/2021.07.19.452930

**Authors:** Dennis M. Echevarria-Cooper, Nicole A. Hawkins, Sunita N. Misra, Alexandra Huffman, Tyler Thaxton, Christopher H. Thompson, Roy Ben-Shalom, Andrew D. Nelson, Anna M. Lipkin, Alfred L. George, Kevin J. Bender, Jennifer A. Kearney

## Abstract

Genetic variants in *SCN2A*, encoding the Na_V_1.2 voltage-gated sodium channel, are associated with a range of neurodevelopmental disorders with overlapping phenotypes. Some variants fit into a framework wherein gain-of-function missense variants that increase neuronal excitability lead to infantile epileptic encephalopathy, while loss-of-function variants that reduce neuronal excitability lead to developmental delay and/or autism spectrum disorder with or without co- morbid seizures. One unique case less easily classified using this binary paradigm is the *de novo* missense variant *SCN2A* p.K1422E, associated with infant-onset developmental delay, infantile spasms, and features of autism spectrum disorder. Prior structure-function studies demonstrated that K1422E substitution alters ion selectivity of Na_V_1.2, conferring Ca^2+^ permeability, lowering overall conductance, and conferring resistance to tetrodotoxin (TTX). Based on heterologous expression of K1422E, we developed a compartmental neuron model that predicted mixed effects on channel function and neuronal activity. We also generated *Scn2a^K1422E^* mice and characterized effects on neurons and neurological/neurobehavioral phenotypes. Dissociated neurons from heterozygous *Scn2a^K1422E/+^* mice exhibited a novel TTX-resistant current with a reversal potential consistent with mixed ion permeation. Cortical slice recordings from *Scn2a^K1442E/+^* tissue demonstrated impaired action potential initiation and larger Ca^2+^ transients at the axon initial segment during the rising phase of the action potential, suggesting mixed effects on channel function. *Scn2a^K1422E/+^* mice exhibited rare spontaneous seizures, interictal EEG abnormalities, altered response to induced seizures, reduced anxiety-like behavior and alterations in olfactory-guided social behavior. Overall, *Scn2a^K1422E/+^* mice present with phenotypes similar yet distinct from *Scn2a* knockout models, consistent with mixed effects of K1422E on Na_V_1.2 channel function.

**Significance Statement:** The early-onset epilepsy variant *SCN2A*-p.K1422E displays unique biophysical properties *in vitro*. To model the impact of this rare variant, we generated *Scn2a^K1422E^* mice. Neurons from heterozygous *Scn2a^K1422E/+^* mice showed functional deficits similar to the loss-of-function effects observed in the *Scn2a* haploinsufficiency model, as well as gain-of-function effects specific to the K1422E variant. There is also some overlap in neurobehavioral phenotypes between *Scn2a^K1422E/+^* and *Scn2a* haploinsufficient mice. However, *Scn2a^K1422E/+^* mice exhibited unique epilepsy-related phenotypes, including epileptiform events and seizures. *Scn2a^K1422E/+^* mice serve as a useful platform to investigate phenotypic complexity of *SCN2A*-associated disorders.

## Introduction

Pathogenic variants in *SCN2A*, which encodes the voltage-gated sodium channel Na_V_1.2 expressed in central nervous system neurons, is a major risk factor for neurodevelopmental disorders (NDD). Within the ion-conducting pore domain of Na_V_1.2 are four critical residues (Asp-Glu-Lys-Ala or DEKA) that confer selectivity for sodium over all other cations (Dudev and Lim, 2014; Sanders et al., 2018) To date, more than 250 heterozygous genetic variants in *SCN2A* have been described in a wide range of NDD (Sanders et al., 2018). Severe gain-of-function (GoF) missense variants, resulting in increased neuronal excitability, lead to infantile epileptic encephalopathy, including Ohtahara syndrome and West syndrome (Ogiwara et al., 2009; Wolff et al., 2017, 2019; Thompson et al., 2020). Less dramatic GoF missense variants lead to benign infantile seizures, which resolve before 2 years-of-age without apparent long-term neurological sequelae (Scalmani et al., 2006; Ben-Shalom et al., 2017). By contrast, loss-of-function (LoF) missense variants and protein-truncating variants (PTVs) that reduce neuronal excitability early in development and synaptic plasticity later in development lead to autism spectrum disorder (ASD) and/or developmental delay (Sanders et al., 2012; Ben-Shalom et al., 2017; Spratt et al., 2019). Some of these cases have co-morbid seizures starting later in life, typically months after onset of encephalopathy associated with GoF variants (Wolff et al., 2017).

While many cases fit neatly into this GoF or LoF paradigm, there are numerous cases that do not, particularly those with seizure onset around one year-of-age (Wolff et al., 2017; Berecki et al., 2018; Sanders et al., 2018). One biophysically remarkable example is the *de novo* missense variant *SCN2A*-p.K1422E, which disrupts one of the four amino acids that constitute the ion selectivity filter (Figure 1A). This variant alters the overall charge of the selectivity filter, converting the positively charged amino acid, lysine (K), to the negatively charged amino acid glutamate (E). The resultant channel, which has been studied in heterologous expression systems, loses sodium selectively, instead becoming a mixed, nonselective cation channel with apparent permeability for sodium, potassium, and calcium with diminished overall conductance (Heinemann et al., 1992; Schlief et al., 1996). Furthermore, the K1422E variant prevents binding of the neuronal sodium channel antagonists tetrodotoxin (TTX) and saxitoxin that bind to the outer vestibule (Terlau et al., 1991; Fozzard and Lipkind, 2010).

**Figure 1.**
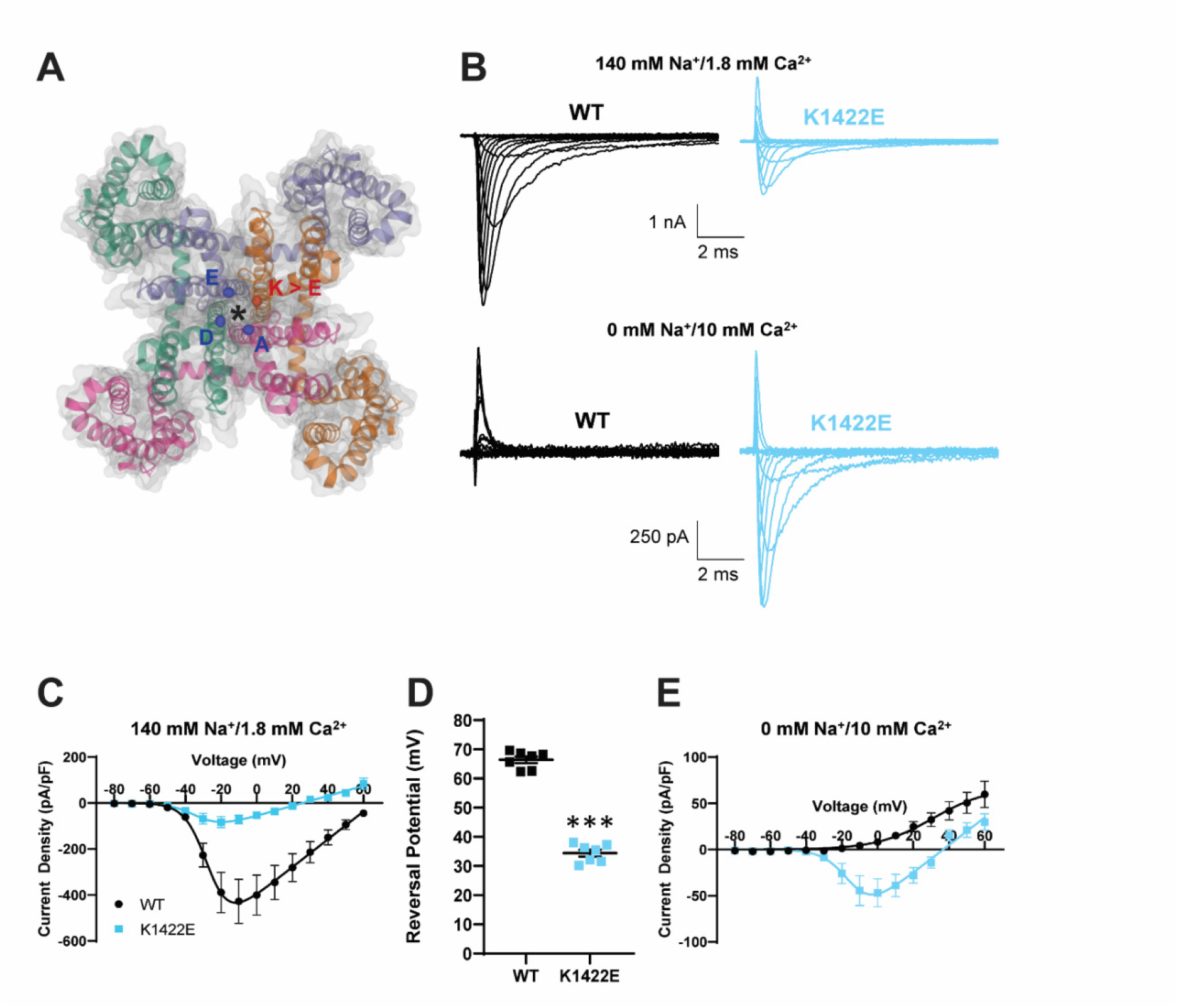
Heterologous expression of K1422E in HEK293T cells reveals altered ion selectivity. **A)** Homology model of voltage gated sodium channel alpha subunit (PDB 6MWA Na_V_Ab, residues1-239). Four internally homologous domains coalesce with four-fold symmetry around an ion conducting pore denoted by an asterisk. The Cα carbons from each residue (DEKA) of the highly conserved ion selectivity filter are represented by colored ellipses. The red ellipse denotes the Cα carbon from the K1422 residue that is substituted for glutamate (E) in the *Scn2a^K1422E^* model. **B)** Representative whole-cell sodium currents (top) and calcium currents (bottom) from WT (left) and K1422E (right) expressing cells. **C)** Summary current-voltage relationship of whole cells sodium current showing reduced sodium current density of K1422E at -10 mV (p=0.0033). **D)** Sodium reversal potential of WT and K1422E expressing cells (p<0.0001). **E)** Summary current-voltage relationship of whole cells calcium current showing increased calcium flux at -10 mV for K1422E (p=0.0109). All data are plotted as mean ± SEM of n=7 cells.

To our knowledge, K1422E is the only variant within the existing *SCN2A* patient population that affects ion selectivity, contrasting markedly with other variants that commonly alter voltage dependence, kinetics, or trafficking (Ben-Shalom et al., 2017; Sanders et al., 2018; Wolff et al., 2019; Adney et al., 2020; Thompson et al., 2020). Given the unique properties of K1422E, it is difficult to hypothesize whether it affects neuronal function in a manner similar to variants that result in gain or loss of function. To investigate this, we generated a mouse model carrying the *SCN2A*-p.K1422E variant and examined effects on cellular and network excitability using electrophysiological and imaging techniques. Excitatory neurons in allocortex and neocortex exhibited features indicative of functional K1422E-containing Na_V_1.2 channels, including the presence of a novel TTX-insensitive current and aberrant calcium influx occurring during the Na_V_- mediated rising phase of the action potential. Analysis of behaving animals revealed a mix of phenotypes, including infrequent spontaneous seizures, reduced anxiety-like behavior, and alterations in olfactory-guided social behavior. Thus, these data suggest that altering Na_V_1.2 ion selectivity results in cellular and behavioral phenotypes that partially mirror those observed in other models of *Scn2a* dysfunction, in addition to features that are entirely unique to K1422E.

## Materials and Methods

### Heterologous cell electrophysiolog

Heterologous expression of human Na_V_1.2 WT (Addgene #162279)(DeKeyser et al., 2021) or K1422E was performed in HEK293T cells. Cells were grown in 5% CO_2_ at 37°C in Dulbecco modified Eagle’s medium supplemented with 10% fetal bovine serum, 2 mM L-glutamine, 50 units/ml penicillin, and 50 μg/ml streptomycin. Cells were transiently co-transfected with Na_V_1.2 and the accessory β_1_ and β_2_ subunits (2 μg total plasmid DNA was transfected with a cDNA ratio of 10:1:1 for Na_V_1.2:β_1_:β_2_ subunits) using Qiagen SuperFect reagent (Qiagen, Valencia, CA, U.S.A.). Human β_1_ and β_2_ cDNAs were cloned into plasmids encoding the CD8 receptor (CD8-IRES-hβ_1_) or enhanced green fluorescent protein (EGFP) (EFGP-IRES-hβ_2_), respectively, as transfection markers, as previously described (Thompson et al., 2011).

Whole-cell voltage-clamp experiments of heterologous cells were performed as previously described (Thompson et al., 2011, 2012). Whole-cell voltage-clamp recordings were made at room temperature using an Axopatch 200B amplifier (Molecular Devices, LLC, Sunnyvale, CA, USA). Patch pipettes were pulled from borosilicate glass capillaries (Harvard Apparatus Ltd., Edenbridge, Kent, UK) with a multistage P-1000 Flaming-Brown micropipette puller (Sutter Instruments Co., San Rafael, CA, USA) and fire-polished using a microforge (Narashige MF-830; Tokyo, JP) to a resistance of 1.5–2.5 MΩ. The pipette solution consisted of (in mM): 10 NaF, 105 CsF, 20 CsCl, 2 EGTA, and 10 HEPES with pH adjusted to 7.35 with CsOH and osmolality adjusted to 300 mOsmol/kg with sucrose. The recording solution was continuously perfused with bath solution containing (in mM): 145 NaCl, 4 KCl, 1.8 CaCl_2_, 1 MgCl_2_, 10 glucose and 10 HEPES with pH adjusted to 7.35 with NaOH and osmolality 310 mOsmol/kg. For sodium-free recordings, the bath solution contained (in mM): 120 NMDG, 4 KCl, 10 CaCl_2_, 2 MgCl_2_, 10 glucose, and 10 HEPES with pH adjusted to 7.35 with HCl and osmolality 310 mOsmol/kg.

### Mice

*Scn2a^K1422E^* mice on the C57BL/6J strain were generated using CRISPR/Cas9 to introduce the modification of K1422 by homology directed repair. A single nucleotide change was introduced in codon 1422 (AAA > GAA), resulting in a glutamate residue being encoded at the modified position. A single guide RNA (TCCTTTAAATGTGGCCTGTA) with low predicted off-target effects, and a 151 bp repair oligo (5’ – CCTTGTTTCCACTTTTACTCTGATAATCTATTTCCTAAACTATAAAAAAGAGAAGAA GTATATATGTTGATTGTTTTACAGGCCACATTT**<u>G</u>**AAGGATGGATGGATATCATGTAT GCAGCTGTTGACTCAAGAAATGTAAGTTTACTTTGG) were delivered to C57BL/6J embryos at the two-cell stage using electroporation by the Northwestern University Transgenic and Targeted Mutagenesis Laboratory.

Potential founders were screened by PCR of genomic DNA using primers outside the repair oligo homology region (Table 1), and the region of interest was cloned into pCR-TOPO (ThermoFisher) and Sanger sequenced. The mosaic K1422E founder was backcrossed to C57BL/6J mice (Jackson Labs, #000664, Bar Harbor, ME) to generate N1 offspring. N1 offspring were genotyped by Sanger sequencing to confirm transmission of the K1422E editing event and were screened for off-target events by Sanger sequencing of all potential sites with <3 mismatches. N1 males with the confirmed on-target event and without predicted off-target events were bred with C57BL/6J females to establish the line *Scn2a^em1Kea^* (MGI:6390565), which is maintained as an isogenic strain on C57BL/6J by continual backcrossing of *Scn2a^K1422E/+^* heterozygous mice (abbreviated *Scn2a^E/+^*) with inbred C57BL/6J mice. Heterozygous *Scn2a^E/+^* and *Scn2a^+/+^* (wild-type, abbreviated WT) mice for experiments were obtained from this breeding at generations >N3.

**Table 1.**
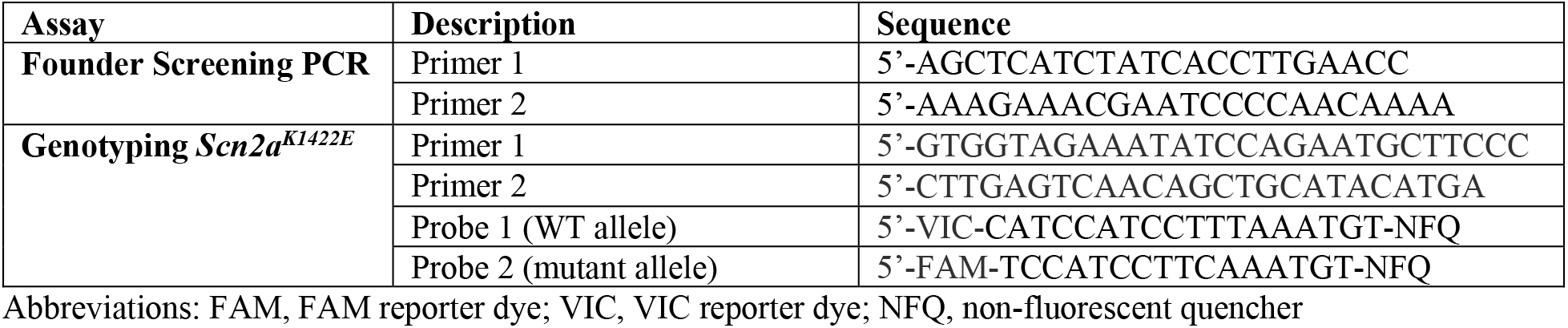
List of Primers and Probes.

Mice were maintained in a Specific Pathogen Free (SPF) barrier facility with a 14-h light/10-h dark cycle and access to food and water ad libitum. Both female and male mice were used for all experiments. All animal care and experimental procedures were approved by the Northwestern University and UC San Francisco Animal Care and Use Committees in accordance with the National Institutes of Health Guide for the Care and Use of Laboratory Animals. Principles outlined in the ARRIVE (Animal Research: Reporting of in vivo Experiments) guideline and Basel declaration (including the 3R concept) were considered when planning experiments.

### Genotyping

DNA was isolated from tail biopsies using the Gentra Puregene Mouse Tail Kit according to the manufacturer’s instructions (Qiagen, Valenica, CA, USA). Genomic DNA was digested with BamHI-HF (New England Biolabs, Ipswich, MA, USA) at 37°C for 1 hour, diluted 1:1 with nuclease-free water and used as template for digital droplet PCR (ddPCR) using ddPCR Supermix for Probes (No dUTP) (Bio-Rad, Hercules, CA, USA) and a custom TaqMan SNP Genotyping Assay (Life Technologies, Carlsbad, CA, USA) (Table 1). Reactions were partitioned into droplets using a QX200 droplet generator (Bio-Rad). PCR conditions were 95°C for 10 min, then 44 cycles of 95°C for 30 s and 60°C for 1 min (ramp rate of 2°C/s) and a final inactivation step of 98 °C for 5 min. Following amplification, droplets were analyzed on a QX200 droplet reader with Quantasoft v1.6.6 software (Bio-Rad).

Whole brain total RNA was extracted from WT and *Scn2a^E/+^* mice at 4 weeks of age using TRIzol reagent according to the manufacturer’s instructions (Invitrogen, Carlsbad, CA, USA). First-strand cDNA was synthesized from 2 µg of total RNA using oligo(dT) primer and Superscript IV reverse transcriptase (RT) according to the manufacturer’s instructions (Life Technologies). First-strand cDNA samples were diluted 1:3 and 5 µL was used as template. Quantitative ddPCR was performed using ddPCR Supermix for Probes (No dUTP) (Bio-Rad) and TaqMan Gene Expression Assays (Life Technologies) for mouse *Scn2a* (FAM-MGB-Mm01270359_m1) and *Tbp* (VIC-MGB-Mm00446971_m1) as a normalization standard. Reactions were partitioned into droplets using a QX200 droplet generator (Bio-Rad).

Thermocycling conditions and analysis were performed as described for genotyping. Both assays lacked detectable signal in no-RT controls. Relative transcript levels were expressed as a ratio of *Scn2a* concentration to *Tbp* concentration (normalized to the mean ratio for WT mice), with 8 biological replicates per genotype, balanced by sex. Statistical comparison between groups was made using Student’s t-test (GraphPad Prism, San Diego, CA, USA).

### Immunoblotting

Whole brain membrane proteins were isolated from WT and *Scn2a^E/+^* mice at 4 weeks of age. Membrane fractions (50 µg/lane) were separated on a 7.5% SDS-PAGE gel and transferred to nitrocellulose. Blots were probed with anti-Na_V_2.1 pAb (Alomone Labs, Jerusalem, ISR; #ASC-002, RRID: AB_2040005; 1:200 dilution) and anti-mortalin/GRP75 mAb (Antibodies Inc, Davis, CA, USA; Neuromab N52A/42, RRID: 2120479; 1:1000 dilution), which served as a normalization standard. Alexa Fluor 790 goat anti-rabbit antibody (Jackson ImmunoResearch, West Grove, PA, USA; #111-655-144, RRID: AB_2338086; 1:10,000 dilution) and 680 goat anti-mouse antibody (Jackson ImmunoResearch; #115-625-146; RRID: AB_2338935; 1:10,000 dilution) were used to detect signal on an Odyssey imaging system (Li-COR, Lincoln, Nebraska USA). Relative protein levels were determined by densitometry with Image Studio (Li-COR) and expressed as a ratio of Na_V_1.2 to GRP75 (normalized to the mean ratio for WT mice), with 8 biological replicates per genotype, balanced by sex. Statistical comparison between groups was made using Student’s t-test (GraphPad Prism).

### Acutely dissociated neuron electrophysiology

Hippocampal pyramidal neurons from P21-P24 WT or *Scn2a^E/+^* were isolated as previously described (Mistry et al., 2014; Thompson et al., 2017). Whole-cell voltage-clamp recordings were made at room temperature using an Axopatch 200B amplifier (Molecular Devices) in the absence and presence of 500 nM tetrodotoxin (TTX). Patch pipettes were pulled from borosilicate glass capillaries (Harvard Apparatus Ltd., Edenbridge, Kent, UK) with a multistage P-97 Flaming-Brown micropipette puller (Sutter Instruments) and fire-polished using a microforge (Narashige MF-830) to a pipette resistance of 1.5–2.5 MΩ. The pipette solution consisted of (in mM) 5 NaF, 105 CsF, 20 CsCl, 2 EGTA, and 10 HEPES with pH adjusted to 7.35 with CsOH and osmolarity adjusted to 280 mOsmol/kg with sucrose. The recording chamber was continuously perfused with a bath solution containing in (mM) 20 NaCl, 100 N-methyl-D-glucamine, 10 HEPES, 1.8 CaCl_2_ꞏ2H_2_O, 2 MgCl_2_ꞏ6H_2_O, and 20 tetraethylammonium chloride with pH adjusted to 7.35 with HCl and osmolarity adjusted to 310 mOsmol/kg with sucrose.

Voltage-clamp pulse generation and data collection were done using Clampex 10.4 (Molecular Devices). Whole-cell capacitance was determined by integrating capacitive transients generated by a voltage step from −120 mV to −110 mV filtered at 100 kHz low pass Bessel filtering. Series resistance was compensated with prediction >70% and correction >90% to assure that the command potential was reached within microseconds with a voltage error <3 mV. Leak currents were subtracted by using an online P/4 procedure. All whole-cell currents were filtered at 5 kHz low pass Bessel filtering and digitized at 50 kHz.

### Modeling

Channel biophysical properties and models were generated as described in Spratt et al., 2010, with APs generated with a 2.1 nA current applied at the soma. A K1422E model was generated by first assuming equal permeabilities for Cs^+^ and K^+^, and a relative permeability for Na^+^ vs K^+^ of 1:0.7, as described in Heinemann et al., 1992.

### Ex vivo whole-cell electrophysiology

Experiments were performed in accordance with guidelines set by the University of California Animal Care and Use Committee. 250 µm-thick coronal slices containing medial prefrontal cortex were obtained from *Scn2a^E/+^* and WT littermate mice of both sexes aged P35-45. Cutting solution contained, in mM: 87 NaCl, 25 NaHCO_3_, 25 glucose, 75 sucrose, 2.5 KCl, 1.25 NaH_2_PO_4_, 0.5 CaCl_2_, and 7 MgCl_2_, bubbled with 5% CO_2_/95% O_2_. After cutting, slices were incubated in the same solution for 30 min at 33°C, then at room temperature until recording. Recording solution contained, in mM: 125 NaCl, 2.5 KCl, 2 CaCl_2_, 1 MgCl_2_, 25 NaHCO_3_, 1.25 NaH_2_PO_4_, and 25 glucose, bubbled with 5% CO_2_/95% O_2_, with an osmolarity of ∼310 mOsm.

For whole-cell recording of layer 5b pyramidal cells, slices were visualized using Dodt contrast optics on a purpose built 2-photon microscope. Patch electrodes were pulled from Schott 8250 glass (3-4 MΩ tip resistance) and filled with a solution containing, in mM: 113 K- gluconate, 9 HEPES, 4.5 MgCl_2_, 14 Tris2-phosphocreatine, 4 Na2-ATP, 0.3 Tris-GTP, 600 μM OGB-5N, 0.1 μM EGTA and 20 μM AlexaFluor 594; ∼290 mOsm, pH 7.2-7.25.

Electrophysiological data were acquired using a Multiclamp 700B amplifier (Molecular Devices) and custom routines in IgorPro (Wavemetrics, Portland, OR, USA). Data were acquired at 50 kHz and Bessel filtered at 20 kHz. Recordings were made using quartz electrode holders to minimize electrode drift within the slice, enabling stable imaging of diffraction-limited spots in close proximity to the recording electrode (Sutter Instruments). Recordings were excluded if series resistance exceeded 14 MΩ or if the series resistance changed by greater than 20% over the course of the experiment. Fast pipette capacitance, as measured in cell-attached voltage clamp mode (typically 10-12 pF), was compensated 50% in current-clamp recordings, and data were corrected for a 12 mV junction potential.

### Two-Photon Imaging

Imaging was performed as described previously (Lipkin et al., 2021). A Coherent Ultra II laser was tuned to 810 nm and epifluorescence and transfluorescence were collected through a 60x, 1.0 NA objective and a 1.4 NA oil immersion condenser, respectively (Olympus). Dichroic mirrors and band-pass filters (575 DCXR, ET525/70 m-2p, ET620/60 m- 2p, Chroma) were used to split fluorescence into red and green channels. HA10770-40 photomultiplier tubes (PMTs, Hamamatsu) selected for >50% quantum efficiency and low dark counts captured green fluorescence (Oregon Green BAPTA 5N). Red fluorescence (AlexaFluor 594) was captured using R9110 PMTs (Hamamatsu).

Fluorescence data were collected in a pointscan configuration, where the laser was parked at single diffraction-limited spots along the AIS membrane. Data were collected over a series of 5 points spanning 2 µm (e.g., separated by 0.5 µm), in regions either 3-5 µm from the axon hillock (proximal AIS) or 28-30 µm from the axon hillock (distal AIS). Each point was imaged at 20 kHz for 25 ms preceding and 100 ms following an AP. Individual points were imaged in a sequence of 2,4,1,3,5, with 2 being the point most proximal to the soma. Individual APs within the set of 5 points were separated by 250 ms. Data were averaged over 40 repetitions for each site and subsequently averaged over all 5 spots for a total averaging of 200 sampled APs. Data were then smoothed using a 40-point binomial filter in IgorPro for analysis. Calcium transients were normalized to saturating conditions (ΔG/G_sat_) as previously described (Lipkin et al., 2021). Transient onset was defined as the time at which signals exceeded root-mean-squared noise levels of the 20 ms period preceding AP onset.

### Video-EEG monitoring

Male and female WT and *Scn2a^E/+^* mice were implanted with prefabricated 3-channel EEG headmounts (Pinnacle Technology, Lawrence, KS, USA) at 4-6 weeks of age. Briefly, mice were anesthetized with ketamine/xylazine and place in a stereotaxic frame. Headmounts with four stainless steel screws that serve as cortical surface electrodes were affixed to the skull with glass ionomer cement. Anterior screw electrodes were 0.5-1 mm anterior to bregma and 1 mm lateral from the midline. Posterior screws were 4.5-5 mm posterior to bregma and 1 mm lateral from the midline. EEG1 represents recordings the right posterior to left posterior (interelectrode distance ≈ 2 mm). EEG2 represents recordings form right anterior to left posterior (interelectrode distance ≈ 5 mm). The left anterior screw served as the ground connection. Following at least 48 h of recovery, tethered EEG and video data were continuously collected from freely moving mice with Sirenia acquisition software (Pinnacle Technology) using a sampling rate of 400 Hz as previously described (Hawkins et al., 2016). At least 96 h of EEG data were acquired from each subject (WT range: 96-672 h/mouse (n = 17 mice; 5-11 weeks of age); *Scn2a^E/+^* range: 168-672 h/mouse (n = 15 mice; 5-13 weeks of age)). Raw data was notch filtered with a 1 Hz window around 60 and 120 Hz prior to analysis. Video-EEG records were manually reviewed with Sirenia software, MATLAB (MathWorks, Natick, MA, USA) and EEGLAB (Swartz Center for Computational Neuroscience, CA, USA) by 2 independent reviewers blinded to genotype. Spontaneous seizures were defined as isolated events with an amplitude of ≥ 3 times baseline, duration of ≥10 s, and that show evolution in power and amplitude. Epileptiform discharges were defined as isolated events with a spike and overriding fast activity, an amplitude of ≥ 3 times baseline, duration of 150-500 ms, and with increased power in frequencies > 20 Hz compared to baseline. Samples with high baseline artifact were excluded from analysis.

### Flurothyl seizure induction

Susceptibility to seizures induced by the chemoconvulsant flurothyl (Bis(2,2,2-trifluoroethyl) ether, Sigma-Aldrich, St. Louis, MO, USA) was assessed in WT and *Scn2a^E/+^* mice at 6-9 weeks of age. Flurothyl was introduced into a clear, plexiglass chamber (2.2 L) by a syringe pump at a rate of 20 µL/min and allowed to volatilize. Latencies to first myoclonic jerk, generalized tonic-clonic seizure (GTCS) with loss of posture, and time interval between these phases were recorded (n = 19-20 per genotype and sex). Groups were compared using Student’s t-test (GraphPad Prism), with sexes considered separately.

### Neurobehavioral assays

Male and female WT and *Scn2a^E/+^* mice were tested between 6 and 11 weeks of age. Male and female mice were tested separately with at least a one-hour delay between sessions. For all experiments, mice were acclimated in the behavior suite with white noise for 1 h prior to behavioral testing. At the end of each procedure, mice were placed into a clean cage with their original littermates. Behavioral testing was performed by experimenters blinded to genotype. For the initial cohort of mice, evaluation occurred over 6 weeks, with one assay performed each week: week 1- zero maze; week 2- light-dark exploration; week 3- open field; week 4- rotarod; week 5- three-chamber social interaction; and week 6- olfactory habituation/dishabituation test. The remaining assays were performed on a new cohort of mice, with one assay performed each week: week 1- marble burying; week 2- Y-Maze (not reported); and week 3- buried food. For all measures, males and females were considered separately. Statistical comparison between groups were made using Student’s t-test or two-way repeated measures ANOVA with Sidak’s post-hoc comparisons, unless otherwise indicated (Table 2).

**Table 2.**
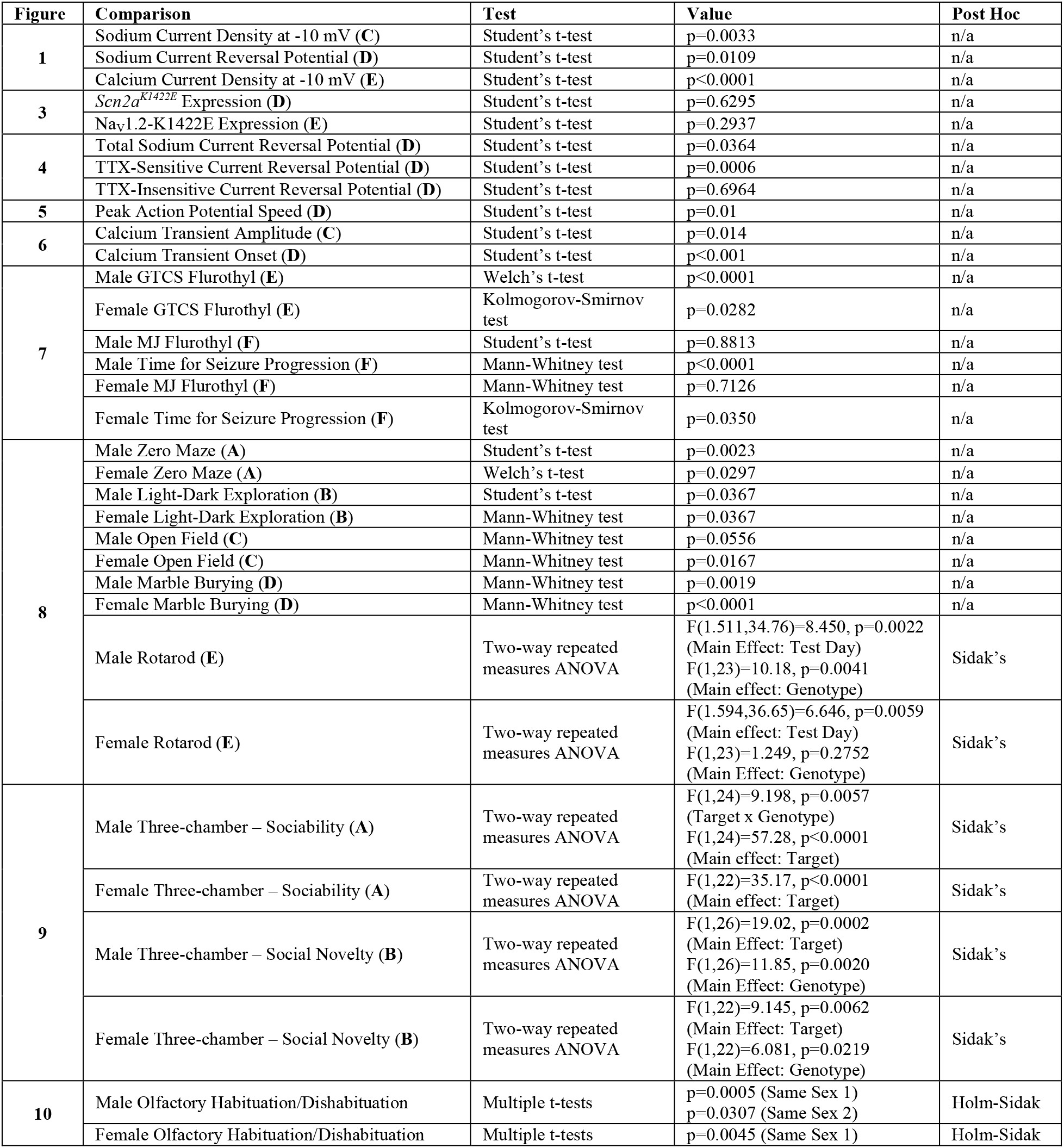
Statistical Comparisons.

### Zero maze

Mice were evaluated for anxiety-related behavior in an elevated zero maze at 6 weeks of age. The maze consists of an annular platform (diameter- 46 cm; elevation- 50 cm) that is divided into equally sized quadrants, alternating between open and enclosed (wall height- 17 cm). This configuration lacks the ambiguous center region associated with the elevated plus maze (Shepherd et al., 1994). Individual mice were placed near an enclosed arm of the maze and allowed to freely explore for 5 min. Limelight software (Actimetrics, Wilmette, IL, USA) was used to video record each trial. Ethovison XT software (Noldus, Leesberg, VA, USA) was used to track the position of the mouse, and calculate distance traveled, mean velocity, and time spent in closed or open arms (n = 12-14 per genotype and sex). Trials where mice fell off the maze were excluded from analysis.

### Light-dark exploration

Mice were evaluated for anxiety-related behavior in a light-dark box at 7 weeks of age. The plexiglass box is divided into equally sized light (400 lx) and dark (0 lx) sections (20 x 40 cm) separated by a central partition plate with a small opening (3 x 5 cm) to allow transit between sections. Individual mice were placed in the center of the light section, facing away from the dark section, and allowed to freely explore for 10 min. Limelight software was used to video record each trial. Ethovison XT software was used to track the position of the mouse, and calculate time spent in light or dark sections (n = 13-15 per genotype and sex).

### Open field

Mice were evaluated for baseline activity and anxiety-related behavior in an open field at 8 weeks of age. Individual mice were placed in the center of an open field arena (46 x 46 cm) and allowed to freely explore for 30 min. Limelight software was used to video record each trial. Ethovison XT software was used to track the position of the mouse, and calculate total distance traveled, mean velocity, and time spent in the periphery (9 cm from wall) and center (28 x 28 cm) (n = 13-15 per genotype and sex).

### Rotarod

Mice were evaluated for motor coordination and balance using an accelerating rotarod (Panlab, Harvard Apparatus, Barcelona, Spain) at 9 weeks of age. Up to 5 mice were placed on the rotating rod that accelerated from 4 to 40 RPM over 5 min. Mice were given three trials per day for 3 consecutive days with an inter-trial interval of 15 min. Latency to fall from the rotating rod was automatically recorded (n = 11-14 per genotype and sex).

### Three-chamber social interaction

Mice were evaluated for sociability and preference for social novelty using a three-chamber social interaction test at 10 weeks of age. The three-chamber apparatus consists of a plexiglass box divided into equally sized chambers (21 x 42 cm) separated by transparent plates with small openings (3 x 5 cm) to allow transit between chambers. All chambers were subject to uniform illumination (170 lx). One wire cylinder-shaped cage was placed in the center of each side chamber and used to enclose a sex-matched, 8–12- week-old C57BL/6J stranger mouse during the testing phase of the assay. During an initial habituation phase, individual test mice were placed in the middle of the center chamber and allowed to freely explore for 10 min. During the first test phase (sociability), a stranger mouse was randomly placed in a wire cage in one of the side chambers and the test mouse was allowed to freely explore for 10 min. During the second testing phase (social novelty), the stranger mouse from the sociability phase remained in place and was then considered familiar. Then, a second stranger mouse was placed in the wire cage in the opposite side chamber and the test mouse was allowed to freely explore for 10 min. Limelight software was used to video record each trial. Ethovison XT software was used to track the position of the test mouse, and calculate time spent in each chamber and time spent sniffing stranger mice or empty wire cage (n = 11-14 per genotype and sex). Trials where mice spent >10 % of trial time on top of the wire cages were excluded.

### Olfactory habituation/dishabituation test

Olfactory impairment could interfere with social behaviors, therefore olfactory discrimination for social and non-social odor was evaluated in 11 week old mice as previously described (Yang and Crawley, 2010). Dry applicators were used to present odor stimuli to test mice: non-social odors- distilled water, almond extract (McCormick, Hunt Valley, MD, USA; 1:100 dilution), and banana extract (McCormick; 1:100 dilution); social odors- unfamiliar social cage (same sex), unfamiliar social cage (opposite sex). During an initial habituation phase, individual mice were placed in a clean testing cage with a clean dry applicator and allowed to freely explore for 30 min. During the testing phase, odor stimuli were presented to test mice for three consecutive trials each (15 trials total) in the following order: (1) distilled water, (2) almond extract, (3) banana extract, (4) same sex cage, (5) opposite sex cage. Each odor was presented for 2 min with an inter-trial interval of 1 min. Trials were video recorded and sniffing time was evaluated manually by an independent reviewer blinded to genotype (n = 13-14 per genotype and sex).

### Marble burying task

Marble burying was evaluated in 6-week-old mice to assess phenotypes related to anxiety- and obsessive-compulsive behavior. Individual mice were placed in a clean testing cage with 4 cm of bedding and acclimated for 15 min. Mice were then briefly removed from the cage while bedding was flattened and 20 marbles were evenly placed across the cage in a 5 x 4 matrix with a small open space at the front of the cage. A baseline image of the cage was taken prior to reintroduction of the mouse and reimaged after the 30-min trial. The two images were compared for the number of marbles buried, defined by at least 50 % of the marble being submerged under the bedding (n = 13-14 per genotype and sex).

### Buried food test

Mice were evaluated for olfactory-guided behavior using a buried food task at 8 weeks of age as previously described (Yang and Crawley, 2010). Teddy Grahams (Nabisco, Hanover, NJ, USA) have been established as a palatable food stimulus in this assay. In order to familiarize test mice with the stimulus odor, one cookie was placed in the home cage of all subjects for 3-4 consecutive days prior to testing. In order to drive olfactory-guided behavior, mice were transferred to a clean cage with their original littermates without access to food 15-18 h before testing. On the testing day, single cookies were buried in a random corner of clean cages with 3 cm of bedding. Individual mice were placed in a clean testing cage and latency to find the food stimulus was recorded (n = 13-14 per genotype and sex). Mice that failed to find the food stimulus after 15 min received a latency score of 900 s.

### Statistical Analysis

Table 2 summarizes statistical tests used for all comparisons along with computed values. D’Agostino & Pearson tests for normality were used to determine parametric versus non-parametric test selection. F test to compare variances was used to determine where to apply correction for unequal standard deviations.

## Results

### K1422E alters channel ion permeability in heterologous expression systems

Previous biophysical characterizations of rat Na_V_1.2 channels expressed in *Xenopus* oocytes suggested that K1422E alters channel ion selectivity, increasing permeation for potassium and calcium (Heinemann et al., 1992; Schlief et al., 1996). Thus, before assessing K1422E function in mice, we first assessed channel function in HEK293T cells. Current density was significantly lower in cells expressing K1422E compared to WT channels (Current density at -10 mV: WT: - 427.6 ± 95.7 pA/pF, n=7, K1422E: -72.0 ± 18.7 pA/pF, n=7, p=0.0033) (Figure 1B-C).

Furthermore, the reversal potential was more hyperpolarized (WT: 66.4 ± 1.1 mV, n=7, K1422E: 34.4 ± 1.2 mV, n=7, p<0.0001; Figure 1D), suggestive of enhanced potassium permeability. To test whether K1422E channels were also permeable to calcium, recordings were made in sodium-free solutions with a higher calcium concentration (10 mM). Consistent with previous studies of bacterial and vertebrate Na_V_1 channels (Favre et al., 1996; Naylor et al., 2016), the WT channel showed no inward calcium-mediated current under these conditions (WT: 4.3 ± 0.5pA/pF, n=7). By contrast, K1422E channels supported appreciable inward current (WT: -44.4 ± 16.2 pA/pF, n=7, p=0.0109), indicating that the variant promoted calcium permeable (Figure 1B, E).

Based on previous biophysical characterizations of K1422E and our own performed here (Heinemann et al., 1992; Schlief et al., 1996), we developed a channel model and compartmental neuronal model to provide testable predictions for how K1422E affects neuronal function (Figure 2). The K1422E variant was modeled first by introducing potassium permeability to an established Na_V_1.2 channel model at a ratio of 1:0.7 (Na:K), and assuming that Cs^+^ and K^+^ permeability were comparable, as determined previously (Heinemann et al., 1992). Subsequently, calcium permeability was increased to levels that best captured the reversal potential observed in HEK293T cells, with reversals for Na^+^, Cs^+^ (≈ K^+^), and Ca^2+^ determined from the Goldman-Hodgkin-Katz equation (Figure 1C). This was best described by relative permeabilities for Na^+^, K^+^ and Ca^2+^ of 1:0.7:0.8. Resultant current density relative to WT was reduced modestly in the model, but less markedly as observed in experiments (Figure 2A, B; Heinemann et al., 1992). This is because calcium has been shown to antagonize K1422E channels, reducing overall current density (Heinemann et al., 1992). We therefore developed an additional channel model with reduced current density to match empirical observations, which was mimicked by reducing the number of K1422E channels relative to WT. This K1422E channel model was incorporated into previously established models of cortical pyramidal cells (Ben-Shalom et al., 2017) at a 50:50 ratio with a wild-type Na_V_1.2 allele, and at reduced relative density to mimic increased calcium antagonism.

**Figure 2.**
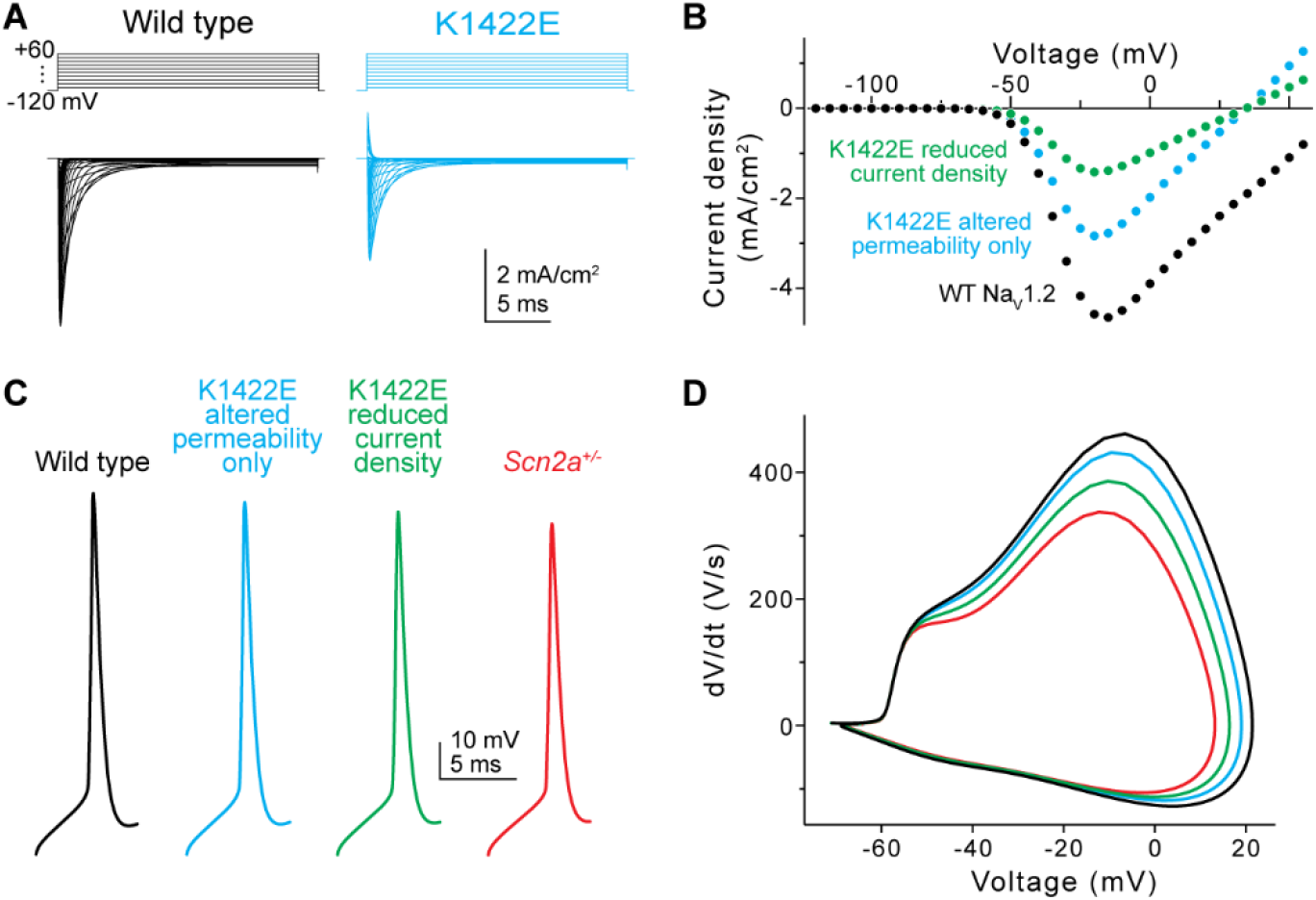
*In silico* simulations of K1422E in neocortical pyramidal cells. **A)** Current evoked from simulated WT (black) and K1422E (cyan) channels in response to voltage steps from -120 to +60 mV (10 mV increments). **B)** Current-voltage relationship for WT and K1422E channels. With altered permeability for K and Ca, K1422E current were reduced ∼50% compared to WT, with a reversal at +33 mV (cyan). Reductions in current density to mimic calcium antagonism were required to match empirical results (green, compare to Figure 1C). **C)** APs measured via a somatic electrode site in a model of a neocortical layer 5b pyramidal cell (identical to Spratt et al., 2019). +/K1422E conditions were modeled with or without reduced current density (cyan, green) and compared to WT (+/-) and haploinsufficient conditions (+/-, red). **D)** Phase-plane plots of APs in C.

APs generated by somatic current injection had features consistent with partial LoF conditions, with reductions in peak AP speed that approached, but did not match those observed in *Scn2a^+/-^* model neurons (Figure 2C-D; 6.4, 16.2, and 26.8% for reduction in AP speed for K1422E without calcium antagonism, with calcium antagonism, and *Scn2a^+/-^* conditions, respectively).

### Generation and initial characterization of Scn2a^K1422E^ Mice

We developed an *in vivo* model of the NDD-associated *SCN2A* p.K1422E pathogenic variant by using CRISPR/Cas9 genome editing to introduce the K1422E single nucleotide variant into mouse *Scn2a* via homology directed repair (Figure 3A). *Scn2a^K1422E/+^* heterozygous mutants (abbreviated as *Scn2a^E/+^*) were born at the expected Mendelian ratios and there was no difference in body weight compared to WT littermates when measured at 4 weeks (WT: 13.5 ± 0.36 g, n=13, *Scn2a^E/+^*: 13.6 ± 0.49 g, n=14, p=0.8349, Student’s t-test). We used droplet digital RT-PCR (RT-ddPCR) and immunoblotting to evaluate whole brain expression of *Scn2a* transcript and Na_V_1.2 protein, respectively, and observed no difference in expression between *Scn2a^E/+^* and WT mice (Figure 3B-D). Similar to constitutive knockout of *Scn2a*, mice homozygous for K1422E (*Scn2a^E/E^*) exhibit 100% mortality by postnatal day 1 (Planells-Cases et al., 2000). Because *SCN2A*- associated NDD is associated with heterozygous variants, our experiments focused on comparing *Scn2a^E/+^* mice to WT littermate controls.

**Figure 3.**
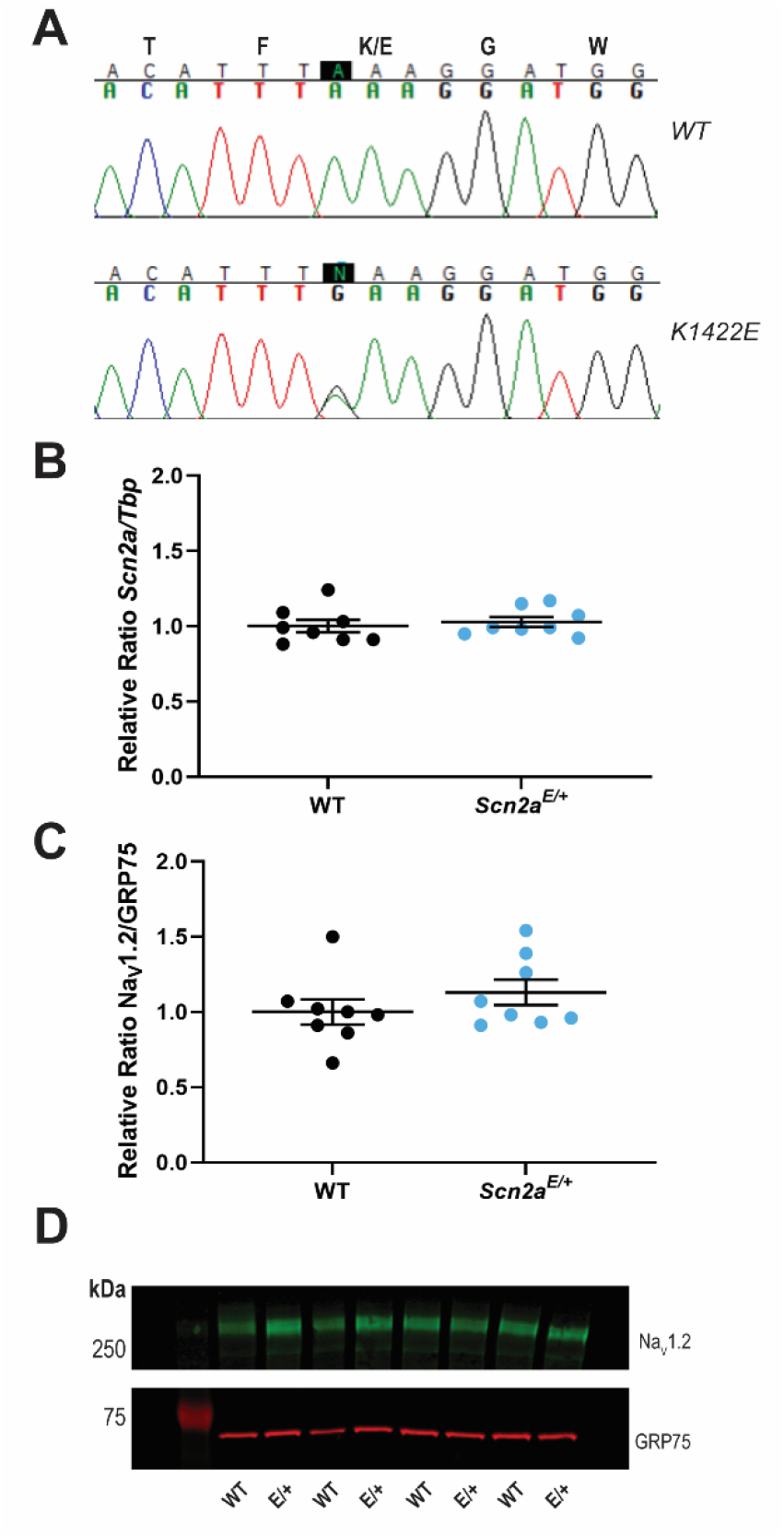
Generation and molecular characterization of *Scn2a^K1422E^* mice. **A)** Sequencing chromatograms of *Scn2a* genomic PCR products with the first nucleotide of the K1422E codon highlighted in black. Top chromatogram from a WT littermate control mouse shows homozygosity for the WT nucleotide at the K1422E codon. Bottom chromatogram from a heterozygous *Scn2a^E/+^* mouse shows heterozygosity for the single nucleotide change introduced by CRISPR/Cas9 genome editing and homology directed repair. **B)** Relative expression of whole brain *Scn2a* transcript in WT and *Scn2a^E/+^* mice assayed by RT-ddPCR. Relative transcript levels are expressed as a ratio of *Scn2a* concentration to *Tbp* concentration (normalized to WT average). There was no difference in transcript expression between genotypes (p=0.6295; n=8 mice per genotype). **C)** Relative expression of whole brain Na_V_1.2 protein in WT and *Scn2a^E/+^* mice assayed by immunoblotting. Quantification is expressed as a ratio of Na_V_1.2 immunofluorescence relative to GRP75/Mortalin (normalized to WT average). There was no difference in protein expression between genotypes (p=0.2937; n=8 mice per genotype). For both **B** and **C**, circles represent samples from individual mice, horizontal lines represent mean, and error bars represent SEM. **D)** Representative immunoblot. Bands corresponding to Na_V_1.2 (MW: 260 kDa) are visualized in green (Alexa Fluor 790) while bands corresponding to GRP75 (MW: 73 kDa) are visualized in red (Alexa Fluor 680).

### Scn2a^K1422E^ channels affect pyramidal cell excitability in allocortex and neocortex

Na_V_1.2 channels are expressed in excitatory pyramidal cells throughout hippocampus and neocortex (Hu et al., 2009; Lorincz and Nusser, 2010). To determine whether K1422E channels are functionally expressed in *Scn2a^E/+^* mice, we performed a series of experiments in dissociated neuronal culture and ex vivo using acute slices. First, we acutely isolated hippocampal neurons from P24-25 WT and *Scn2a^E/+^* mice and assessed sodium currents derived from perisomatic channels maintained in the cells. This includes somatic sodium channels, which are predominantly Na_V_1.6 isoforms in hippocampal neurons (Lorincz and Nusser, 2010; Spratt et al., 2019), but also a fraction of the proximal axon initial segment, which is enriched with Na_V_1.2 channels (Lorincz and Nusser, 2010). Consistent with current arising largely from Na_V_1.6 isoforms, dissociated cell sodium current were not altered appreciably in *Scn2a^E/+^* conditions; however, the reversal potential of this sodium current was hyperpolarized ∼6 mV in *Scn2a^E/+^* cells (WT: 38.5 ± 2.2 mV n=11, *Scn2a^E/+^*: 32.7 ± 1.0 mV, n=9, p=0.0364; Figure 4A, D). This suggests that K1422E channels may be contributing to this current, albeit at levels that cannot be resolved within cell-to-cell variability in total current amplitude. To further resolve K1422E channels in such preparations, we reassessed currents in 500 nM TTX, as the K1422E mutation has been shown to reduce TTX sensitivity (Terlau et al., 1991). In these conditions, *Scn2a^E/+^* neurons produced an appreciable TTX-resistant current that reversed at more hyperpolarized potentials (WT: 35.6 ± 3.9 mV n=8, *Scn2a^E/+^*: 16.5 ± 2.7 mV, n=9, p=0.0006; Figure 4B, D). Importantly, the remaining TTX-sensitive currents, which is not contaminated by currents mediated by K1422E-containing channels, were comparable to WT in current density and reversal potential (WT: 43.9 ± 1.7 mV n=10, *Scn2a^E/+^*: 46.8 ± 2.9 mV, n=9, p=0.0364; Figure 4C-D).

**Figure 4.**
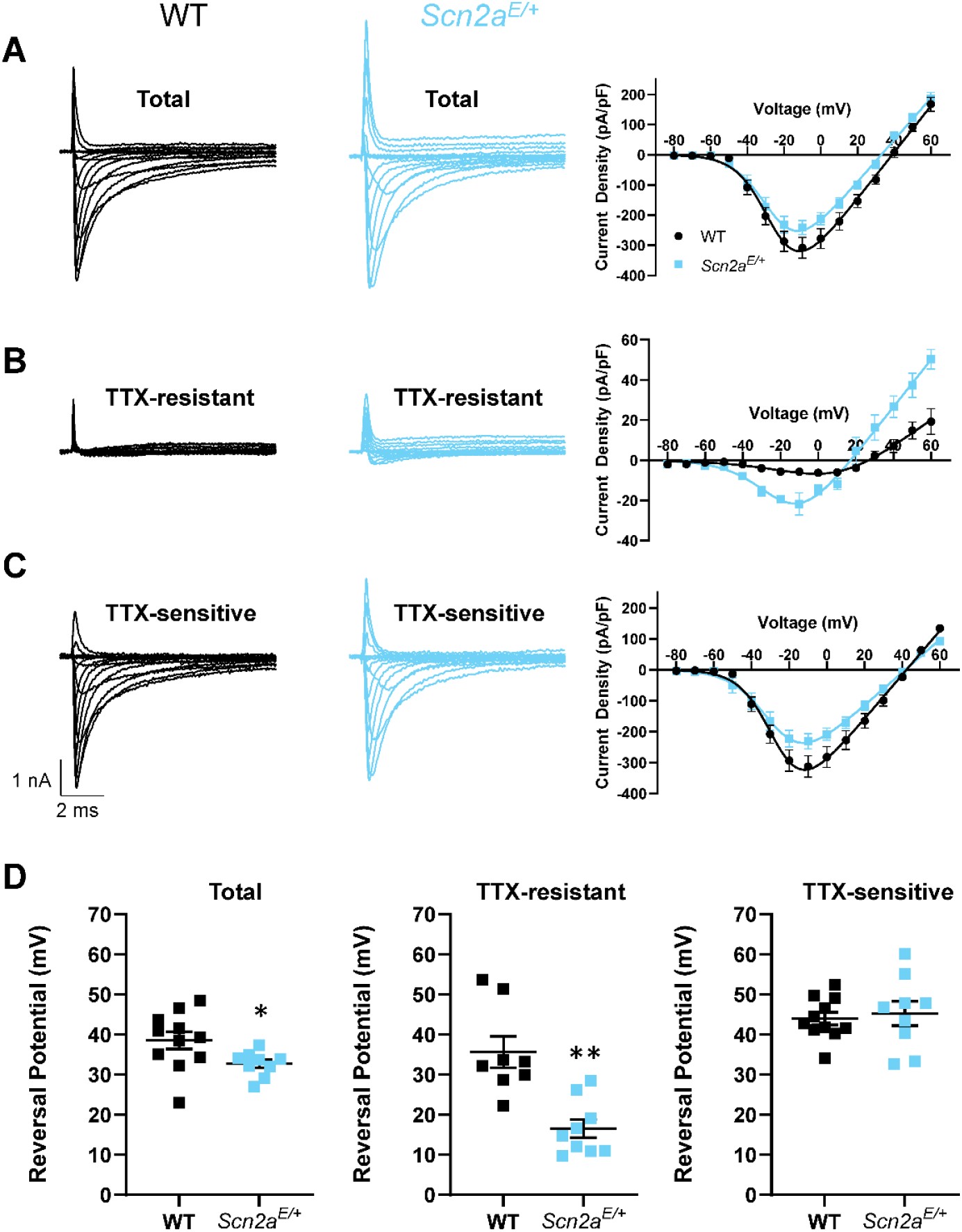
Whole-cell sodium currents of acutely isolated neuron. Representative whole-cell sodium currents and current voltage relationships of **A)** total sodium current, **B)** TTX-resistant currents, and **C)** TTX-sensitive currents from acutely dissociated hippocampal pyramidal neurons from WT and *Scn2a^E/+^* mice, **D)** Sodium reversal potential of total sodium current (left, p=0.0364), TTX-resistant current (middle, p=0.0006), and TTX-sensitive currents (right, p=0.6964). All data are plotted as mean ± SEM of n=8-11 cells.

Na_V_1.2 channels contribute to a larger fraction of somatic sodium influx in neocortical pyramidal cells. In these cells, progressive loss of *Scn2a* alleles, either from constitutive heterozygous knockout, or conditional homozygous knockout, leads to progressive decrements in the speed of the somatic component of an AP (Spratt et al., 2019, 2021). Based on the reduced current density observed for K1422E above, we hypothesized that the AP waveform would be similarly affected by this variant. To test this, we made whole-cell current-clamp recordings from layer 5b pyramidal cells in acute slices containing medial prefrontal cortex of *Scn2a^E/+^* mice aged P37-45. *Scn2a^E/+^* pyramidal cells had slower peak AP speed (WT: 605 ± 18 V/s, n = 14; E/+: 527 ± 21 V/s, n = 13; p = 0.01, unpaired t-test), but were otherwise indistinguishable from cells assayed from littermate controls (Figure 5). This reduction in AP speed (13%) is smaller than that observed in *Scn2a^+/-^* heterozygotes (27%; Spratt et al. 2021), consistent with a reduction, but not elimination, of current density through channels with the K1422E variant.

**Figure 5.**
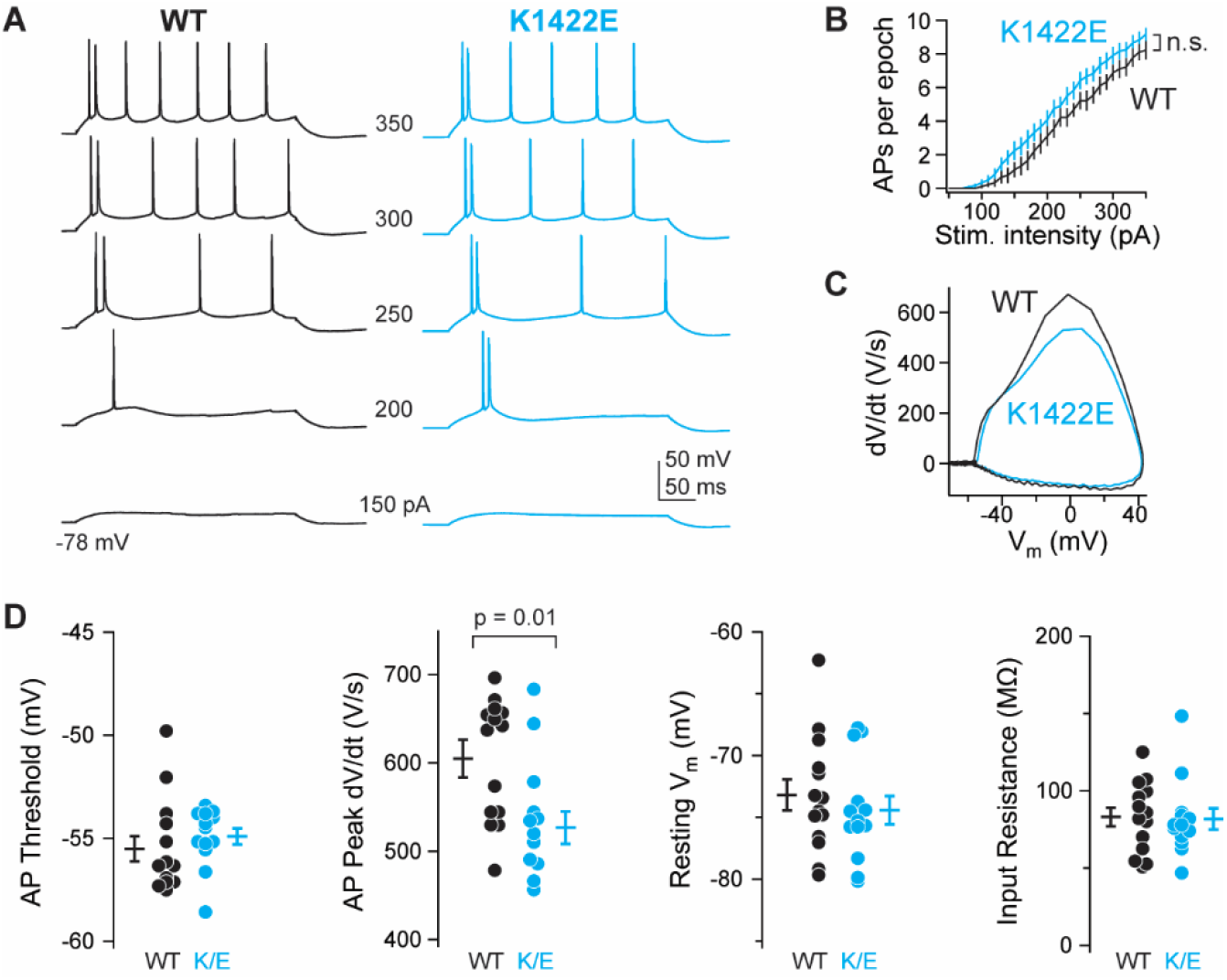
*Scn2a^K1422E^* prefrontal pyramidal cell AP waveform has loss-of-function characteristics. **A)** Example whole-cell voltage response to somatic current injection in WT (black) and *Scn2a^E/+^* (K1422E, cyan) neurons. Numbers between examples correspond to current injection amplitude. **B)** AP number per 300 ms current injection, color coded as in A. Bars are mean ± SEM. n = 13 cells each. **C)** Rheobase AP as dV/dt vs. voltage (phase-plane plot). Note reduction of peak dV/dt for *Scn2a^E/+^* cells compared to WT, indicative of loss-of-function in Na_V_1.2-mediated somatic depolarization. **D)** Summaries of AP waveform and intrinsic excitability characteristics. Circles are single cells, bars are mean ± SEM. Peak dV/dt is reduced in *Scn2a^E/+^* cells, unpaired t-test. n = 14 WT, 13 *Scn2a^E/+^* cells.

Furthermore, changes in AP speed were best fit by compartmental model predictions that combined changes in channel permeabilities with a reduction in current density due to calcium antagonism (predicted 16% reduction) (Figure 2).

These changes in AP waveform suggest that K1422E-Na_V_1.2 channels are functional in neocortical pyramidal cells. To test whether calcium influx through Na_V_1.2 channels can be observed in intact neurons, we imaged AP-evoked calcium transients in the AIS with high spatiotemporal precision using 2-photon pointscan imaging. Under normal conditions, calcium influx occurs during AP repolarization in all regions of the axon, including the AIS, and can be separated temporally from sodium influx occurring during AP depolarization (Geiger and Jonas 2000; Ritzau-Jost et al. 2014; Rowan, Tranquil, and Christie 2014; Filipis and Canepari 2021; Lipkin et al. 2021; but see Hanemaaijer et al. 2020). But in cells expressing K1422E channels, calcium influx may also occur during AP depolarization in regions enriched with Na_V_1.2 channels. In mature pyramidal neurons, Na_V_1.2 channels are clustered densely in a region of the AIS proximal to the soma, whereas Na_V_1.6 channels are enriched in the distal AIS (Figure 6A; Hu et al., 2009). Therefore, we imaged the proximal and distal initial segment, 5 µm and 30 µm from the axon hillock, respectively, corresponding to regions enriched or lacking Na_V_1.2 channels (Figure 6B). Consistent with a lack of Na_V_1.2 channels in the distal AIS, there were no differences in the amplitude or timing of AP-evoked calcium transients in the distal AIS. By contrast, calcium transients were larger and occurred earlier, before AP membrane potential peaked, in the proximal AIS of *Scn2a^E/+^* cells (Amplitude, WT: 2.8 ± 0.2 ΔG/G_sat_, E/+: 3.8 ± 0.3 ΔG/G_sat_,p = 0.014; timing relative to AP peak, WT: 0.62 ± 0.15 ms, n = 12, E/+: -0.17 ± 0.13 ms, n = 10; p < 0.001, unpaired t-test; Figure 6B-D). Thus, these data indicate that Na_V_1.2-K1422E channels flux calcium into the cell, and that this influx occurs during AP depolarization in the AIS.

**Figure 6.**
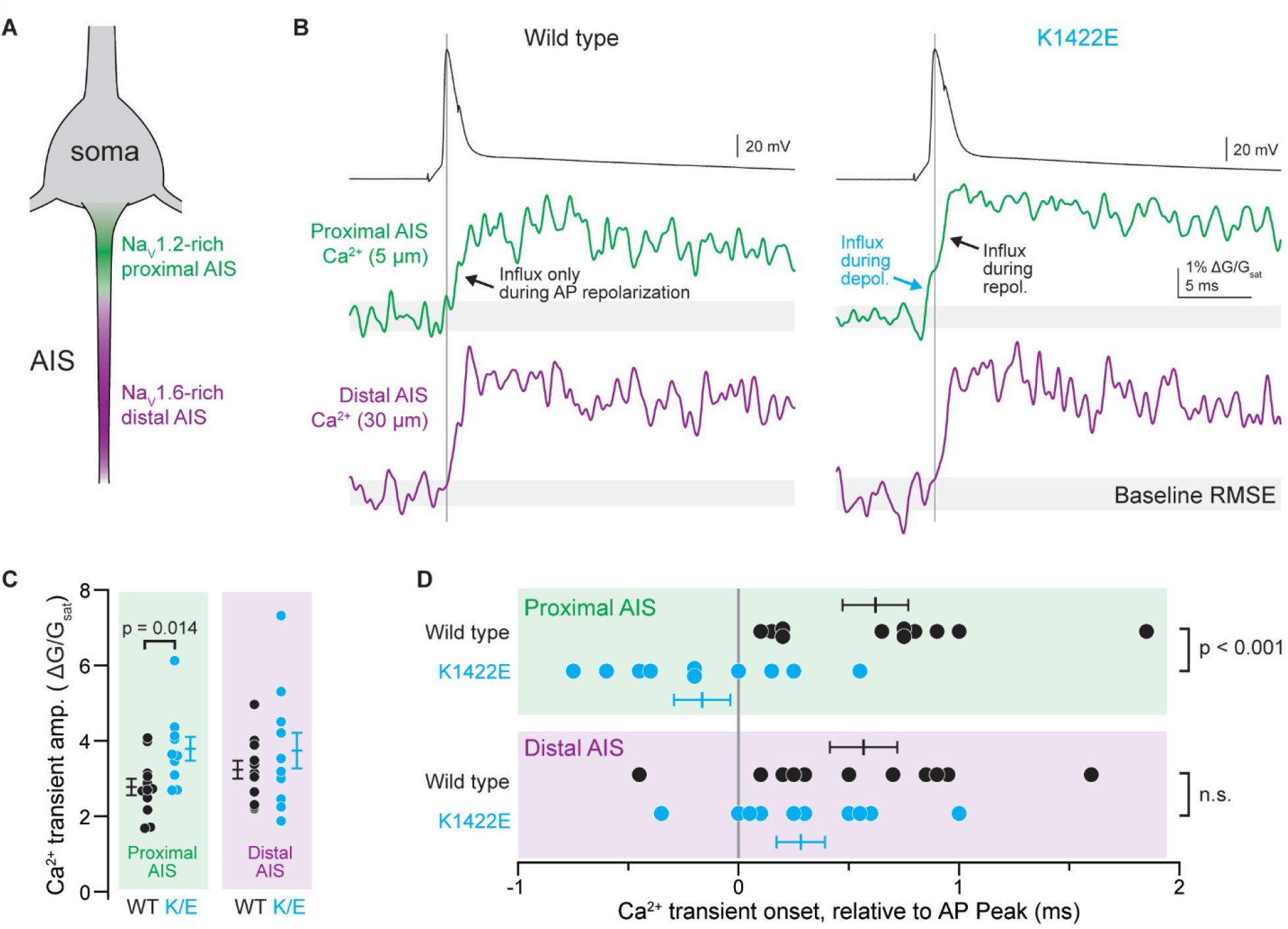
AP-evoked Ca^2+^ influx during the rising phase of the AP in the proximal AIS of *Scn2a^E/+^* cells. **A)** Pyramidal cell initial segments are enriched with Na_V_1.2 proximal to the soma and Na_V_1.6 more distal to the soma. Pointscan imaging was performed 5 and 30 µm from the axon hillock, corresponding to Na_V_1.2 and Na_V_1.6-enriched regions, respectively. **B)** Examples of AP-evoked (2 nA, 2 ms stimulus; black, top) calcium transients imaged in pointscan mode in the proximal (green, middle) and distal (violet, bottom) AIS in WT (left) and *Scn2a^E/+^* cells (right). Vertical line is aligned to peak AP voltage. Grey shaded area encompasses imaging signal root-mean-squared error (RMSE) during baseline, before AP. Consistent deviation above this error value defines onset of Ca^2+^ transient. Note Ca^2+^ influx before AP peak in proximal AIS of K1422E condition, only. **C)** Amplitude of Ca^2+^ transient is higher in proximal AIS of *Scn2a^E/+^* cells, consistent with influx from both local voltage-gated calcium channels and additional influx through K1422E Na_V_1.2 channels. Circles are single cells and bars are mean ± SEM. p values from unpaired t-tests. **D)** Ca^2+^ transient onset occurs earlier in the proximal AIS of *Scn2a^E/+^* cells, consistent with Ca^2+^ influx through K1422E Na_V_1.2 channels. Display as in **C**.

### Scn2a^K1422E^ mice exhibit abnormalities in EEG and alterations in seizure threshold

The child with the *SCN2A* p.K1422E variant initially presented with treatment-refractory infantile spasms at 13 months of age and went on to develop other seizure types (Sundaram et al., 2013). As previously discussed, seizure phenotypes have been associated with GoF and LoF effects on *SCN2A,* in both humans and mice (Ogiwara et al., 2009, 2018; Gazina et al., 2010; Sanders et al., 2018). The above data suggest that K1422E has mixed effects on channel function: namely, LoF-like effects with respect to neuronal excitability, as well as aberrant ion influx. How this mixed channel phenotype might affect brain activity and contribute to seizure susceptibility is unknown. Therefore, to evaluate *Scn2a^E/+^* mice for spontaneous seizures and epileptiform events, juvenile (4-6 weeks) mice were implanted with EEG headmounts for video-EEG monitoring which occurred from 5-11 weeks of age. *Scn2a^E/+^* mice exhibited spontaneous localized seizures that occurred during sleep without any observable behavioral changes (Figure 7A-B). These seizures occurred rarely (< 1 per week) and were not observed in all *Scn2a^E/+^* mice. Similar events were never observed in WT mice. *Scn2a^E/+^* mice also exhibited interictal epileptiform discharges, including isolated high amplitude spikes with overriding fast activity that had higher power across low and high frequencies up to 170 Hz (Figure 7C-D). These events occurred at a rate of 2-5 events per 8 hours when quantified across a window that included 4-hour epochs from the light and dark phases.

**Figure 7.**
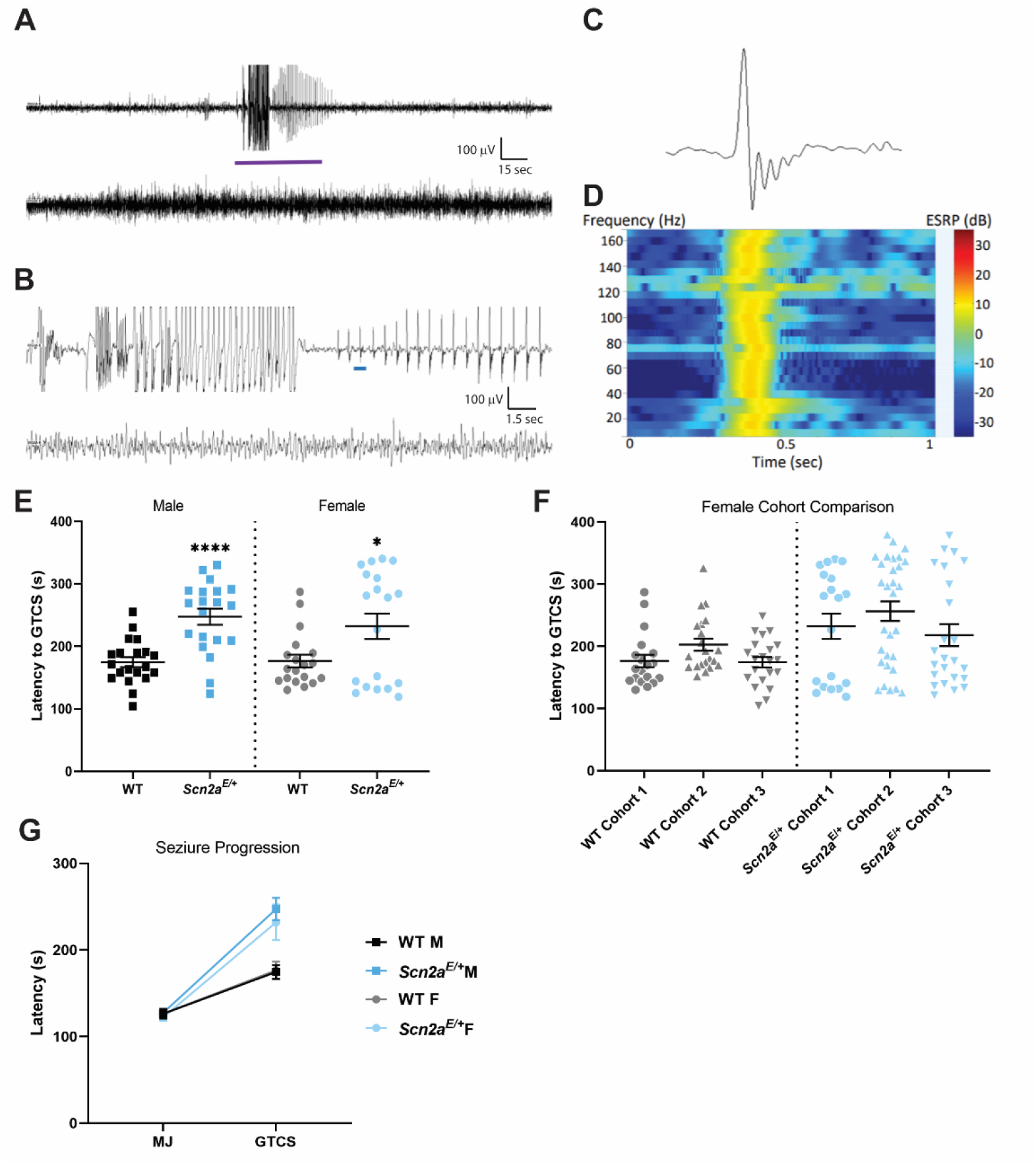
EEG abnormalities and altered susceptibility to induced seizures in *Scn2a^E/+^* mice. **A)** Representative 5-minute epoch of EEG from *Scn2a^E/+^* mice. A localized seizure occurred as an abrupt onset of rhythmic 2 Hz sharp waves with overriding fast activity that evolve in amplitude and frequency for ∼45 seconds before abruptly terminating with return to typical sleep background. **B)** 30-second epoch corresponding to the purple bar segment from **A**. The top line in both **A** and **B** corresponds to channel 1 (right posterior-left posterior) and second line is channel 2 (right anterior-left posterior). **C)** Example of an isolated high amplitude sharp wave with overriding fast activity corresponding to the blue bar segment in **B**. **D)** Power spectrum for the sharp wave in **C** showing elevated power in decibels across the 1-170 Hz frequency range at the time of discharge. **E)** Latency to flurothyl-induced GTCS in *Scn2a^E/+^* mice compared to WT at 6-9 weeks of age. *Scn2a^E/+^* males had an elevated threshold for flurothyl-induced seizures compared to WT (WT: 175 ± 8 sec, *Scn2a^E/+^*: 247 ± 13 sec, ****p<0.0001; Welch’s t-test). *Scn2a^E/+^* females also had an elevated threshold for flurothyl-induced seizures compared to WT (WT: 177 ± 10 sec, *Scn2a^E/+^*: 232 ± 20 sec, *p=0.0282; Kolmogorov-Smirnov test). Symbols represent samples from individual mice, horizontal lines represent mean, and error bars represent SEM. **F)** Latency to first flurothyl-induced generalized tonic-clonic seizure (GTCS) in WT and *Scn2a^E+^* female mice across multiple cohorts (n = 19-30 per genotype and cohort). Cohorts were evaluated at different times. Data from *Scn2a^E/+^* females is abnormally distributed in all three cohorts compared to cohort-matched WT controls. **G)** Average latency to first myoclonic jerk (MJ) and GTCS, with connecting line depicting time of progression between stages. There was no genotype difference in latency to first MJ for both sexes (Table 2). However, progression between stages was slower for both male and female *Scn2a^E/+^* mice compared to WT (Males: p<0.0001; Mann-Whitney test; Females: p=0.0350, Kolmogorov-Smirnov test). Symbols represent group mean and error bars represent SEM; n=19-20/sex/genotype Males and females were analyzed separately.

Since these spontaneous seizures in *Scn2a^E/+^* mice appear to be rare and difficult to observe due to a lack of an obvious behavioral component, we next asked how the *Scn2a^K1422E^* variant affects seizure susceptibility in an induced-seizure paradigm. We used the volatile chemoconvulsant flurothyl, a GABA_A_ antagonist (Ferland, 2017), to induce a stereotyped progression to generalized tonic-clonic seizures (GTCS) in juvenile (6-9 weeks of age) *Scn2a^E/+^* and WT mice. While latency to first myoclonic jerk was not affected by genotype, latency to first GTCS (defined as characteristic limb flexion/extension with loss of posture) was affected by genotype (Table 2). Both male and female *Scn2a^E/+^* mice had a higher threshold for flurothyl-induced seizures compared to sex-matched WT littermates (p < 0.0001 for males; p = 0.0282 for females). Average latency was 247 ± 13 sec for *Scn2a^E/+^* males, 175 ± 8 sec for WT males, 257 ± 21 sec for *Scn2a^E/+^* females, and 177 ± 10 sec for WT females (Figure 7E). We also noted that the data reflecting latency to first GTCS in *Scn2a^E/+^* females was abnormally distributed compared to data from WT females. This effect was reproducible in three separate cohorts of *Scn2a^E/+^* and WT females (Figure 7F). Time to progress from the first myoclonic jerk to the first GTCS was also affected by genotype (Table 2). Both males and female *Scn2a^E/+^* mice progressed more slowly compared to sex-matched WT controls (p < 0.0001 for males and p = 0.035 for females). Average progression time was 120 ± 13 sec for *Scn2a^E/+^* males versus 48 ± 8 sec for WT males, and 109 ± 19 sec for *Scn2a^E/+^* females versus 52 ± 10 sec for WT females (Figure 7G). The above data suggest that rare spontaneous seizures occur in *Scn2a^E/+^* mice, but they remain localized in parieto-occipital cortex. Furthermore, slower progression to flurothyl-induced GTCS indicates resistance to seizure spreading in *Scn2a^E/+^* mice, consistent with the localized nature of spontaneous seizures observed using EEG.

### Lower anxiety-related behavior in Scn2a^K1422E^ mice

There is evidence to suggest that individuals with ASD are at increased risk for co-morbid anxiety (van Steensel et al., 2011; Perihan et al., 2021) and anxiety-related behavior is frequently assessed in mouse models of ASD-related genes, including *Scn2a* (Silverman et al., 2010; Léna and Mantegazza, 2019; Spratt et al., 2019; Tatsukawa et al., 2019). We used zero maze, light-dark exploration, and open field assays to assess anxiety-related behavior in *Scn2a^E/+^* and WT mice. These assays take advantage of mouse thigmotaxis and phototaxic aversion to define anxiety-related behavior. More time spent in exposed, well-lit areas compared to dark, enclosed areas indicates lower anxiety-related behavior. In the zero-maze assay, time spent in the open versus closed arms of the maze was significantly higher in both male and female *Scn2a^E/+^* mice compared to sex-matched WT controls (p = 0.0023 for males; p = 0.0297 for females, respectively). On average, *Scn2a^E/+^* males spent nearly twice as long (30.5 ± 4.1%) in the open arms compared to WT males (15.3 ± 2.2%; Figure 8A). *Scn2a^E/+^* females also spent more time in the open arms (34.8 ± 4.9%) compared to WT females (22.0 ± 2.0%; Figure 8A). During the light-dark exploration assay, time spent in the exposed light zone versus the enclosed dark zone was significantly affected by genotype (p=0.0367 for both male and female comparisons). On average, *Scn2a^E/+^* males spent more time (29.2 ± 3.2%) in the light zone compared to WT males (20.9 ± 2.0%; Figure 8B). *Scn2a^E/+^* females also spent more time in the light zone (32.1 ± 3.5%) compared to WT females (21.1 ± 1.7%; Figure 8B). In the open field assay, time spent in the exposed center zone of the apparatus versus the periphery was significantly affected by genotype when comparing *Scn2a^E/+^* and WT females (p=0.0167), but not when comparing males (p=0.0556). However, the overall genotype effect recapitulated what was seen in the zero maze and light-dark exploration assays, with *Scn2a^E/+^* females spending an average of 10.1 ± 1.2% of test time in the center zone and WT females averaging 6.27 ± 0.9% of time (Figure 8C).

**Figure 8.**
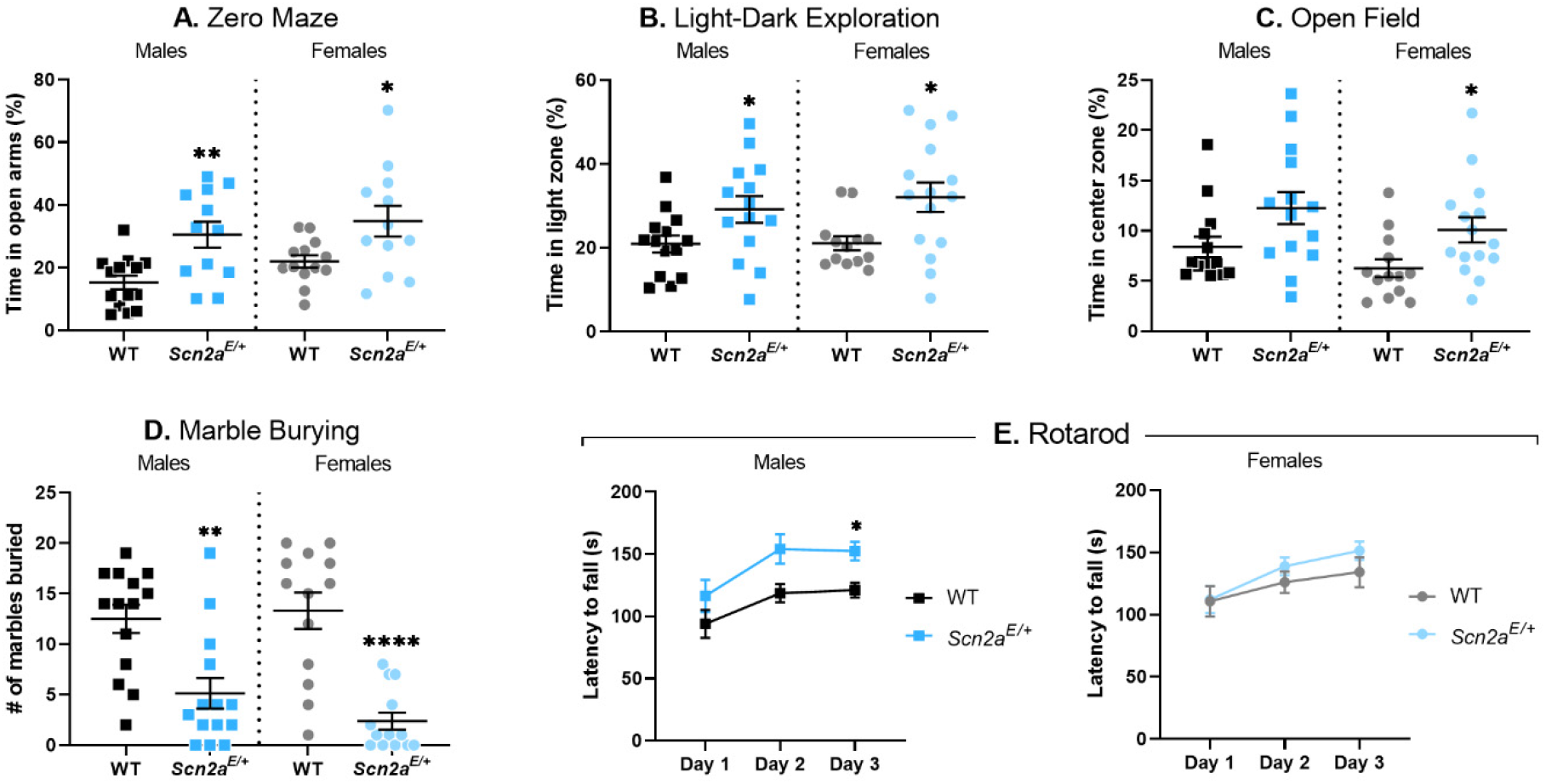
Altered anxiety-related behavior and rotarod performance in *Scn2a^E/+^* mice. **A)** Percent time spent in the open arms of a zero-maze apparatus in *Scn2a^E/+^* mice compared to WT at 6 weeks of age. *Scn2a^E/+^* males spent significantly more time in the open arms compared to WT (WT: 15.3 ± 2.2%, *Scn2a^E/+^*: 30.5 ± 4.1%, **p=0.0023; Student’s t-test). *Scn2a^E/+^* females spent significantly more time in the open arms compared to WT (WT: 22.0 ± 2.0%, *Scn2a^E/+^*: 34.8 ± 4.9%, *p=0.0297; Welch’s t test). **B)** Percent time spent in the light zone of a light/dark box in *Scn2a^E/+^* mice compared to WT at 7 weeks of age. *Scn2a^E/+^* males spent significantly more time in the light zone compared to WT (WT: 20.9 ± 2.0%, *Scn2a^E/+^*: 29.2 ± 3.2%, *p=0.0367; Student’s t-test). *Scn2a^E/+^* females also spent significantly more time in the light zone compared to WT (WT: 21.1 ± 1.7%, *Scn2a^E/+^*: 32.1 ± 3.5%, *p=0.0367; Mann-Whitney test). **C)** Percent time spent in the center zone of an open field apparatus in *Scn2a^E/+^* mice compared to WT at 8 weeks of age. There was not a significant difference in the amount of time spent in the center zone between *Scn2a^E/+^* and WT males (p=0.0556, Mann-Whitney test). However, *Scn2a^E/+^* females spent significantly more time in the center zone compared to WT (WT: 6.27 ± 0.9%, *Scn2a^E/+^*: 10.1 ± 1.2%, *p=0.0167; Mann-Whitney test). **D)** Number of marbles buried during a 30-minute trial by *Scn2a^E/+^* mice compared to WT at 6 weeks of age is displayed. *Scn2a^E/+^* males buried significantly fewer marbles compared to WT (WT: 12 ± 1, *Scn2a^E/+^*: 5 ± 2, **p=0.0019; Mann-Whitney test). *Scn2a^E/+^* females also buried significantly fewer marbles compared to WT (WT: 13 ± 2, *Scn2a^E/+^*: 2 ± 1, ****p=0.0001; Mann-Whitney test). **E)** Average latency to fall during an accelerating rotarod task measured on three consecutive days in *Scn2a^E/+^* mice compared to WT at 9 weeks of age. Daily performance for each animal was assessed by averaging across three trials. Two-way repeated measures ANOVA comparing average latency to fall between *Scn2a^E/+^* and WT males showed significant main effects of test day [*F*(1.511,34.76) = 8.450, **p=0.0022] and genotype [*F*(1,23) = 10.18, *p=0.0041]. *Scn2a^E/+^* males took significantly longer to fall compared to WT on day 3 (WT: 121 ± 6 sec, *Scn2a^E/+^*: 152 ± 7 sec, p=0.0106; Sidak’s post-hoc test). Two-way repeated measures ANOVA comparing average latency to fall between *Scn2a^E/+^* and WT females showed a significant main effect of test day only [*F*(1.594,36.65) = 6.646, **p=0.0059]. For panels **A-D** symbols represent individual mice, horizontal lines represent mean, and error bars represent SEM. For panel **E** symbols and error bars represent mean ± SEM. Males and females were analyzed separately, with n=12-14 per genotype for males and n=11-15 per genotype for females.

Data from our initial set of behavioral assays suggested that *Scn2a^E/+^* mice showed lower anxiety-like behavior compared to WT controls. To further assess these behavioral abnormalities in the context of a more specific behavior involving novel objects, we used a marble burying assay. Both *Scn2a^E/+^* males and females buried significantly fewer marbles compared to WT controls (males, WT: 13 ± 1, *Scn2a^E/+^*: 5 ± 2, p = 0.0019; females, WT: 13 ± 2, *Scn2a^E/+^*: 2 ± 1, p < 0.0001; Figure 8D). Early on, it became apparent that some animals were not burying the marbles and a limited number of subsequent trials were video recorded. We observed normal exploratory behavior in recordings where animals did not bury the marbles, suggesting that the observed genotype effect on marble burying was due to lower anxiety-like behavior rather than inactivity.

### Enhanced rotarod performance in Scn2a^K1422E^ male mice

Movement disorders have been reported in some children with *SCN2A-*associated infantile epileptic encephalopathy (Sanders et al., 2018) and deficits in motor function could potentially confound results from other behavioral assays. Therefore, we used an accelerating rotarod assay to evaluate motor coordination and balance in *Scn2a^E/+^* mice compared to WT controls at 9 weeks of age. For each subject, latency to fall was measured for three trials and daily performance was assessed by averaging across trials. Two-way repeated measures ANOVA showed significant main effects of test day and genotype when comparing average latency to fall between *Scn2a^E/+^* males and WT controls (Table 2). *Scn2a^E/+^* males spent significantly more time on the rotarod compared to WT males by the third day of testing (p = 0.0106). On day three of testing, average latency to fall was 121 ± 6 sec for WT males and 152 ± 7 sec for *Scn2a^E/+^* males (Figure 8E). Two-way repeated measures ANOVA only showed a significant main effect of test day when comparing *Scn2a^E/+^* and WT females (Table 2). These data indicate that basic motor function in *Scn2a^E/+^* remains intact.

### Altered social behavior in Scn2a^K1422E^ mice

As previously noted, ASD or features of ASD are frequently reported in children with *SCN2A* variants, including SCN2A-p.K1422E (Sundaram et al., 2013). Social behavior deficits are a common feature of ASD (American Psychiatric Association, 2013) and have been extensively studied in animal models of ASD-related gene disruptions (Silverman et al., 2010; Barak and Feng, 2016). We used a three-chamber assay, which takes advantage of innate preferences for novel social interactions, to evaluate social behavior in *Scn2a^E/+^* mice compared to WT controls at 10 weeks of age. During the sociability phase, time spent sniffing either a novel object (empty wire cup) or an unfamiliar mouse was measured (Figure 9A). Two-way ANOVA comparing average sniffing time between *Scn2a^E/+^* males and WT controls during the sociability phase showed a significant main effect of target and a significant interaction between target and genotype (Table 2). On average, *Scn2a^E/+^* males spent significantly more time (132.6 ± 8.3 sec) sniffing an unfamiliar mouse compared to WT (93.3 ± 12.7 sec; Figure 9A). Two-way ANOVA showed only a significant main effect of target when comparing average sniffing time between *Scn2a^E/+^* females and WT controls during the sociability phase (Table 2). These data suggest that preference for social interactions is intact for both male and female *Scn2a^E/+^* mice, while *Scn2a^E/+^* males, but not females, display a greater preference for social interactions compared to WT. During the social novelty phase, time spent sniffing either a familiar mouse or an unfamiliar mouse was measured (Figure 9B). Two-way ANOVA showed significant main effects of target and genotype when comparing average sniffing time between *Scn2a^E/+^* and WT males during the social novelty phase (Table 2). On average, *Scn2a^E/+^* males spent significantly more time (77.1 ± 5.7 sec) sniffing an unfamiliar mouse compared to WT males (48.8 ± 8.7 sec; Figure 9B). Two-way ANOVA also showed significant main effects of target and genotype when comparing average sniffing time between *Scn2a^E/+^* females and WT controls during the social novelty phase (Table 2). On average, *Scn2a^E/+^* females spent 61.9 ± 7.7 sec sniffing an unfamiliar mouse, while WT females spent 41.6 ± 4.5 sec sniffing an unfamiliar mouse (Figure 9B). These data suggest that preference for social novelty is intact for both male and female *Scn2a^E/+^* mice, and that *Scn2a^E/+^* males, but not females, display a greater preference for social novelty compared to WT.

**Figure 9.**
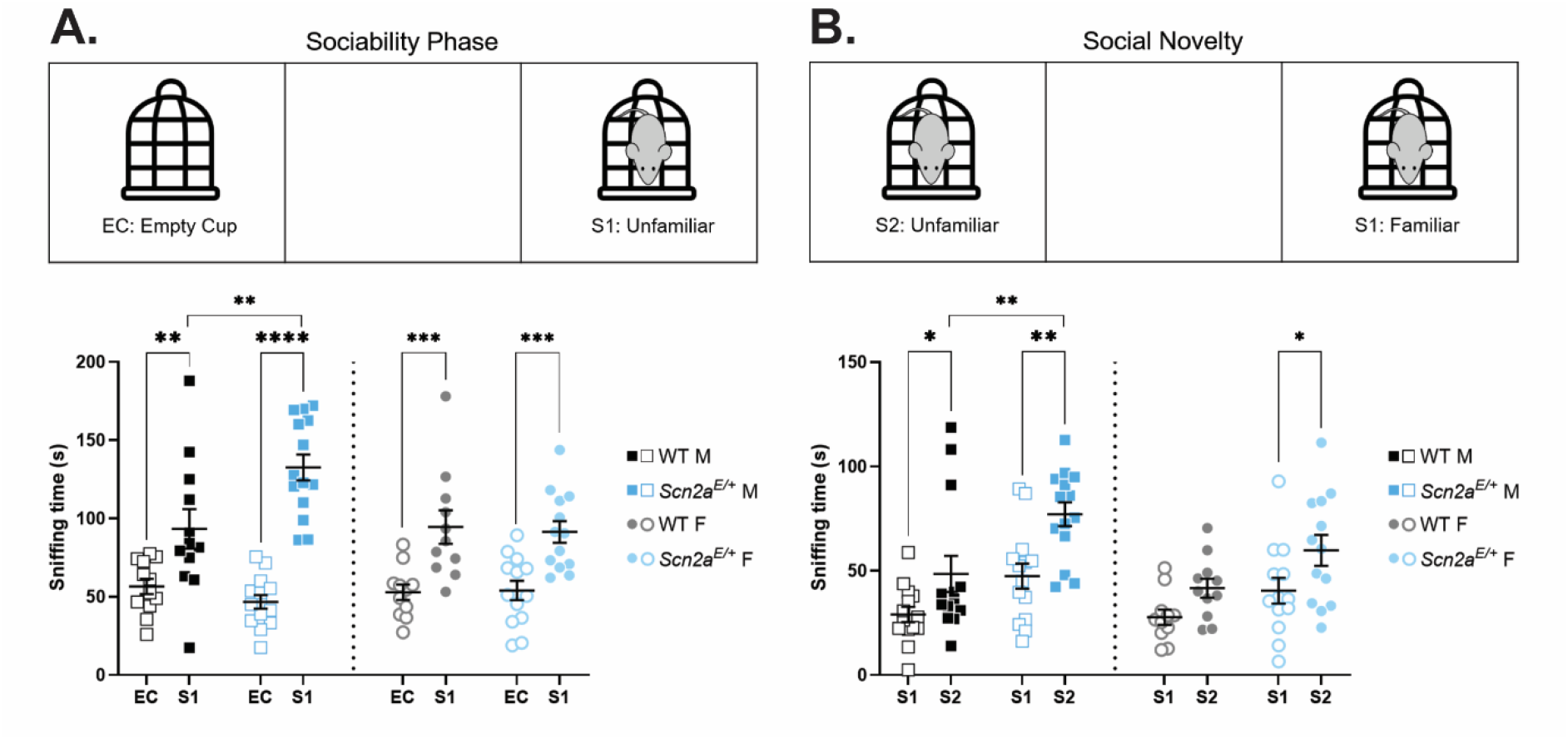
Altered social behavior in *Scn2a^E/+^* mice. **A)** Sociability phase of three-chamber assay. Amount of time spent sniffing either an empty cup (EC) or unfamiliar mouse (S1) in *Scn2a^E/+^* mice compared to WT at 10 weeks of age. Two-way ANOVA (using target as a within-subject variable) comparing average sniffing time between *Scn2a^E/+^* and WT males showed a significant main effect of target [*F*(1,24) = 57.28, p<0.0001] and a significant interaction between target and genotype [*F*(1,24) = 9.198, p=0.0057]. Both *Scn2a^E/+^* and WT males spent significantly more time sniffing an unfamiliar mouse compared to an empty cup (*Scn2a^E/+^*: ****p<0.0001, WT: **p=0.0100; Sidak’s post-hoc test). However, *Scn2a^E/+^* males spent significantly more time sniffing an unfamiliar mouse compared to WT males (WT: 93.3 ± 12.7 sec, *Scn2a^E/+^*: 132.6 ± 8.3 sec, **p=0.0024; Sidak’s post-hoc test). Two-way ANOVA comparing average sniffing time between *Scn2a^E/+^* and WT females showed a significant main effect of target only [*F*(1,22) = 35.17, p<0.0001]. Both *Scn2a^E/+^* and WT females spent significantly more time sniffing an unfamiliar mouse compared to an empty cup (*Scn2a^E/+^*: ***p<0.0007, WT: ***p<0.0008; Sidak’s post-hoc test). **B)** Social novelty phase of three-chamber assay. Amount of time spent sniffing either a familiar mouse (S1) or an unfamiliar mouse (S2) in *Scn2a^E/+^* mice compared to WT at 10 weeks of age. Two-way ANOVA (using target as a within-subject variable) comparing average sniffing time between *Scn2a^E/+^* and WT males showed significant main effects of target [*F*(1,26) = 19.02, p=0.0002] and genotype [*F*(1,21) = 11.85, p=0.0020]. Both *Scn2a^E/+^* and WT males spent significantly more time sniffing an unfamiliar mouse compared to a familiar mouse (*Scn2a^E/+^*: **p=0.0019, WT: *p=0.0432; Sidak’s post-hoc test). However, *Scn2a^E/+^* males spent significantly more time sniffing an unfamiliar mouse compared to WT males (WT: 48.5 ± 8.7 sec, *Scn2a^E/+^*: 77.1 ± 5.7.3 sec, **p=0.0042; Sidak’s post-hoc test). Two-way ANOVA comparing average sniffing time between *Scn2a^E/+^* and WT females also showed significant main effects of target [*F*(1,22) = 9.145, p=0.0062] and genotype [*F*(1,22) = 6.081, p=0.0219]. However, only *Scn2a^E/+^* females spent significantly more time sniffing an unfamiliar mouse compared to a familiar mouse (*Scn2a^E/+^*: *p=0.0327, WT: p=0.1889; Sidak’s post-hoc test). Symbols represent individual mice, horizontal lines represent mean, and error bars represent SEM. Males and females were analyzed separately, with n=12-14 per genotype for males and n=11-13 per genotype for females.

### Lower olfactory dishabituation to social odors in Scn2a^K1422E^ mice

The three-chamber assay described above measures sniffing time as the variable of interest and could therefore be affected by deficits in olfactory discrimination. We used an olfactory habituation/dishabituation assay to evaluate olfactory discrimination in *Scn2a^E/+^* mice compared to WT controls at 11 weeks of age. Each odorant (water, almond extract, banana extract, same sex urine, opposite sex urine) was presented for three consecutive trials before proceeding to the next odor in the sequence for a total of 15 trials. Time spent sniffing the odor delivery apparatus was measured for each trial and compared between genotypes. Habituation refers to a decrease in sniffing behavior upon repeated presentation of an odor, while dishabituation refers to an increase in sniffing behavior upon presentation of a novel odor. Qualitatively, olfactory discrimination and preference for social odors appear to be intact for both male and female *Scn2a^E/+^* mice (Figure 10A). However, the amount of time *Scn2a^E/+^* males spent sniffing same sex urine during the first two presentations was over 50% lower than WT males (p=0.0005 and p=0.0307, respectively).

**Figure 10.**
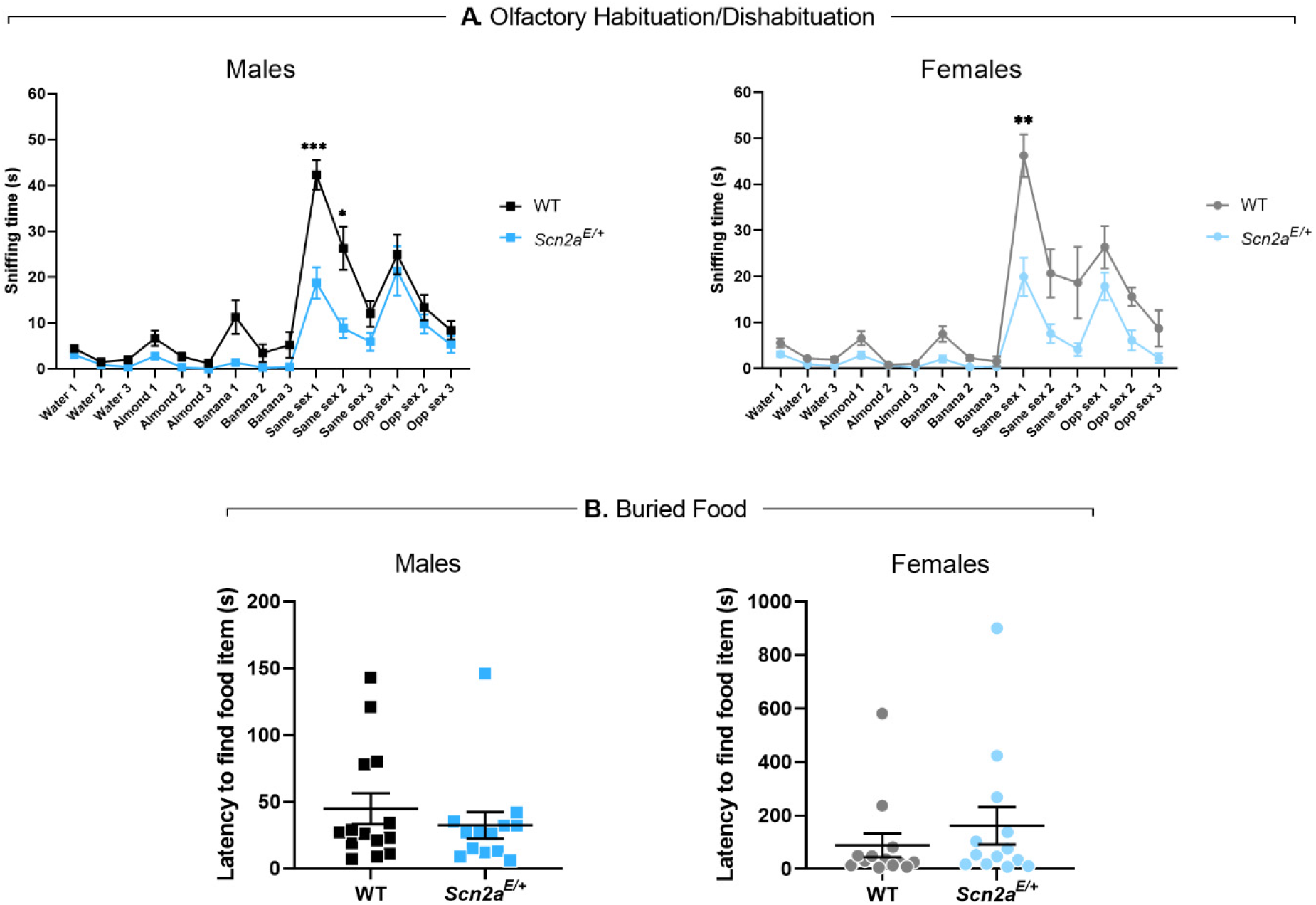
Lower olfactory dishabituation to social odors and intact olfactory-guided behavior in *Scn2a^E/+^* mice. **A)** Average sniffing times during an odor habituation/dishabituation assay in *Scn2a^E/+^* mice compared to WT at 11 weeks of age. Overall, olfactory discrimination in *Scn2a^E/+^* males was not significantly different from WT males. However, *Scn2a^E/+^* males spent significantly less time sniffing same sex urine during the first two presentations compared to WT males. During the first presentation, average sniffing time was 42.3 ± 3.3 sec for WT males and 18.8 ± 3.4 sec for *Scn2a^E/+^* males (***p=0.0005, multiple t-tests). During the second presentation, average sniffing time was 26.3 ± 8.9 sec for WT males and 8.9 ± 2.1 sec for *Scn2a^E/+^* males (*p=0.0307, multiple t-tests). Olfactory discrimination in *Scn2a^E/+^* females was also not significantly different from WT females. However, *Scn2a^E/+^* females spent significantly less time sniffing same sex urine during the first presentation compared to WT females (WT: 46.3 ± 4.6 sec, *Scn2a^E/+^*: 19.9 ± 4.2 sec, **p=0.0045, multiple t-tests). Symbols and error bars represent mean ± SEM. Males and females were analyzed separately, with n=14 per genotype for males and n=12-13 per genotype for females. **B)** Latency to find a buried food item in *Scn2a^E/+^* mice compared to WT at 8 weeks of age. Latency to find a buried food item was not significantly different between *Scn2a^E/+^* and WT males (WT: 44.86 ± 11.58 sec, *Scn2a^E/+^*: 32.46 ± 9.95 sec, p = 0.7471; Mann-Whitney test). Latency to find a buried food item was not significantly different between *Scn2a^E/+^* and WT females (WT: 89.00 ± 44.31 sec, *Scn2a^E/+^*: 161.7 ± 70.02, p = 0.2927; Mann-Whitney test). Symbols represent measured values from individual mice, horizontal lines represent mean, and error bars represent SEM. Males and females were analyzed separately, with n=13-14 per genotype for males and n=13 per genotype for females.

During the first presentation, average sniffing time was 42.3 ± 3.3 sec for WT males and 18.8 ± 3.4 sec for *Scn2a^E/+^* males, while average sniffing time for the second presentation was 26.3 ± 8.9 sec for WT males and 8.9 ± 2.1 sec for *Scn2a^E/+^* males (Figure 10A). *Scn2a^E/+^* females also spent significantly less time sniffing same sex urine compared to WT females, but only during the first presentation (p=0.0045). Average sniffing time was 46.3 ± 4.6 sec for WT females and 19.9 ± 4.2 sec for *Scn2a^E/+^* females (Figure 10A).

The three-chamber and olfactory habituation/dishabituation assays suggest that olfactory- guided behavior is altered in *Scn2a^E/+^* mice and that these effects were more pronounced for social odors. In order to exclude the possibility that olfaction was globally affected, we used a buried food task to compare a separate cohort of *Scn2a^E/+^* mice and WT controls at 9 weeks of age. Task performance was evaluated by measuring the amount of time it took for food-deprived subjects to locate a hidden food stimulus. Both males and female *Scn2a^E/+^* mice performed as well as sex-matched WT controls (males, WT: 44.86 ± 11.58 sec, *Scn2a^E/+^*: 32.46 ± 9.95 sec, p = 0.7471; females, WT: 89.00 ± 44.31 sec, *Scn2a^E/+^*: 161.7 ± 70.02, p = 0.2927; Figure 10B), indicating that olfaction remains intact.

## Discussion

*SCN2A* variants have been associated with a wide range of NDD that reflect a complex spectrum of phenotypes (Sanders et al., 2018). Significant attention has been given to the far ends of this phenotypic spectrum, establishing a framework in which *SCN2A* missense variants that result in GoF effects are associated with infantile epileptic encephalopathy, while PTVs that result in LoF effects are associated with ID/ASD that sometimes present with co-morbid seizures starting later in life (Shi et al., 2012; Brunklaus et al., 2014; Ben-Shalom et al., 2017; Wolff et al., 2017; Begemann et al., 2019). More recently, attempts have been made to further refine the genotype- phenotype relationships of *SCN2A*-related disorders (Crawford et al., 2021), an effort that can be supported by animal models of unusual variants. Here, we focused on the variant *SCN2A*- p.K1422E, which we and others have shown alters ion channel selectivity in heterologous expression systems (Heinemann et al., 1992; Schlief et al., 1996). We further characterized cellular and behavioral phenotypes associated with this unique variant in a newly generated mouse model (*Scn2a^K1422E^*). Excitatory neurons in allocortex and neocortex from *Scn2a^E/+^* mice displayed a novel TTX-insensitive current and aberrant calcium influx that occurs during the rising phase of the AP localized to Na_V_1.2-rich regions of the AIS, indicating that the variant channel is functionally expressed in these cells. *Scn2a^E/+^* mice also display neurological/neurobehavioral phenotypes, including infrequent spontaneous seizures, lower anxiety-like behavior, and alterations in olfactory-guided behavior. These phenotypes are similar yet distinct from those observed in other models of *Scn2a* dysfunction, consistent with altered Na_V_1.2 ion selectivity having mixed effects on channel function.

### Effects of altered Na_V_1.2 ion selectivity on neuronal function

Vertebrate voltage-gated sodium channels contain four highly conserved residues (DEKA) that confer selectivity for sodium (Dudev and Lim, 2014; Sanders et al., 2018). These channels evolved from a primordial channel with mixed selectivity, where the selectivity filter has a glutamic acid (E) substituted for lysine (K) in the 3^rd^ transmembrane domain (Zhou et al., 2004; Stephens et al., 2015). Thus, the K1422E variant can be seen as an evolutionary reversion at the selectivity filter (Gur Barzilai et al., 2012; Zakon, 2012). Consistent with previous work on rat Na_V_1.2 channels in *Xenopus* oocytes (Heinemann et al., 1992), we show that substitution of glutamic acid for lysine at position 1422 (DEKA to DEEA) in human Na_V_1.2 confers calcium permeability not evident in wild-type human Na_V_1.2 channels (Figure 1B).

Based on modeling of biophysical data here and in previous reports (Heinemann et al., 1992), we estimated relative permeabilities of K1422E channels to be Na^+^, K^+^, and Ca^2+^ to be 1:0.7:0.8. Interestingly, some invertebrates express more primitive Na_V_1 channels containing the DEKA selectivity filter that nevertheless appear to have some calcium permeability (Meves and Vogel, 1973). This suggests that additional aspects of Na_V_1 evolution, including changes to other residues lining the pore (Kawai et al., 2021), may confer additional ion selectivity/permeability properties. Consistent with this, recurrent *SCN2A* missense variants that affect pore-adjacent arginine residues R397 and R973 eliminate permeation altogether (Ben-Shalom et al., 2017). Future functional studies should therefore include measures of cation permeability in addition to standard kinetic and voltage dependence assays when considering variants affecting the pore domain.

Before the existence of sodium channels, cellular depolarization was mediated by calcium-selective channels or nonselective cation channels (Gur Barzilai et al., 2012; Zakon, 2012). As such, changes in membrane potential were linked to alterations in intracellular calcium and associated downstream calcium-dependent signaling. Sodium channels are thought to have evolved in part to allow cells to separate changes in voltage from calcium signaling. What effects, then, could arise from recombining these processes in Na_V_1.2 channels? Within the AIS, we show that calcium influx occurs both before and after the peak of the AP in K1422E- expressing neurons (Figure 5B-D). This contrasts with typical conditions where sodium and calcium influx are separated on the rising and falling phases of the AP, respectively (Filipis and Canepari, 2021; Lipkin et al., 2021, but see Hanemaaijer et al., 2020). This additional calcium influx could have myriad effects on AIS function, as many AIS components are regulated directly by calcium or through calcium/calmodulin interactions (Wen and Levitan, 2002; Kim et al., 2004; Swain et al., 2015; Clarkson et al., 2017). Additionally, excess calcium influx could affect cellular processes beyond the AIS as Na_V_1.2 channels are also expressed in the somatodendritic domain and throughout the axons of unmyelinated neurons (Gong et al., 1999; Vacher et al., 2008; Lorincz and Nusser, 2010; Zhu et al., 2020). During high-frequency activity, intracellular sodium concentrations can exceed 50 mM in some nerve terminals (Zhu et al., 2020). Excess calcium influx through K1422E channels could therefore affect transmitter release and short-term presynaptic plasticity (Stanley, 2016; Burke and Bender, 2019). Na_V_1.2 channels also influence dendritic excitability in neocortex (Spratt et al., 2019). Thus, excess calcium may affect dendritic integration/plasticity, depending on the location of these channels relative to synaptic inputs.

### Neurological and neurobehavioral phenotypes associated with the K1422E variant

*Scn2a* knockout models have been used to study phenotypes associated with LoF effects on Na_V_1.2 (Léna and Mantegazza, 2019; Spratt et al., 2019; Tatsukawa et al., 2019; Indumathy et al., 2021). As previously discussed, some properties of *Scn2a^E/+^* neurons are similar to those observed in Na_V_1.2 knockouts (reduced AP speed), while others are unique (calcium flux). Correlated with these complex channel properties are neurological/neurobehavioral phenotypes in *Scn2a^E/+^* mice that are similarly mixed.

EEG recording has been used to evaluate neurological phenotypes in *Scn2a^+/-^* mice with conflicting results. Ogiwara and colleagues (2018) described spike-and-wave discharges characteristic of absence epilepsy in *Scn2a^+/-^* mice, while other groups reported no observable seizures in *Scn2a^+/-^* mice (Mishra et al., 2017). However, this may be due to differences in age and/or strain that can affect spike-and-wave discharges and seizure susceptibility (Bergren et al., 2005; Connor et al., 2005; Martin et al., 2007; Bessaih et al., 2012; Frankel et al., 2014; Kehrl et al., 2014; Miller et al., 2014; Calhoun et al., 2016). We performed video-EEG recordings in *Scn2a^E/+^* mice for a minimum of 96 hours per animal, far longer than what has been reported for *Scn2a* knockouts, and observed rare spontaneous seizures localized to parieto-occipital cortex. To date, spontaneous seizures have not been described in any *Scn2a* knockout models.

In order to further probe seizure susceptibility in *Scn2a^E/+^* mice, we induced seizures using flurothyl. Although latency to the first seizure stage of myoclonic jerk did not differ from WT, *Scn2a^E/+^* mice had longer latency to GTCS. This suggests that threshold for seizure generation is not altered, but that *Scn2a^E/+^* mice may be resistant to seizure spread, which would be consistent with the localized spontaneous seizures. Paradoxical seizure thresholds have been observed in other mouse epilepsy models (Amador et al., 2020; Sah et al., 2020) and may reflect the mechanism by which flurothyl, a GABA_A_ antagonist, induces seizures (Ferland, 2017). Interestingly, we also observed a reproducible non-unimodal distribution of GTCS latencies in *Scn2a^E/+^* females compared to WT. One possible explanation is a genotype-dependent interaction with fluctuating sex hormones and/or neurosteroids, which have previously been linked to altered seizure susceptibility in rodents (Woolley, 2000; Scharfman et al., 2005; Gangisetty and Reddy, 2010; Kight and McCarthy, 2014; Joshi and Kapur, 2019; Li et al., 2020).

Behavioral abnormalities reported in *Scn2a^+/-^* mice vary across research groups and may reflect differences in methodological practices (e.g. age or background strain), as well as inherent variability of behavioral data (Anon, 2009; Saré et al., 2021). We observed lower anxiety-like behavior in *Scn2a^E/+^* mice across three exploration assays (zero maze, light-dark exploration, and open field) and an active marble burying task. This is similar to what has been reported in *Scn2a*^+/-^ mice (Léna and Mantegazza, 2019; Spratt et al., 2019; Tatsukawa et al., 2019). *Scn2a^E/+^* mice also displayed hyper-social behavior in a three-chamber assay, while inconsistent effects on social behavior have been reported in *Scn2a^+/-^* mice (Tatsukawa et al., 2019; Indumathy et al., 2021). Although ASD is typically associated with social behavior deficits (Barak and Feng, 2016), hyper-social behavior has been reported in other ASD-associated genetic mouse models (Chao et al., 2010; Mejias et al., 2011). This highlights the heterogeneity of ASD-associated phenotypes and complexity of modeling human social behavior in animals. In particular, mouse social behavior is guided by olfaction and olfactory function must be considered when interpreting assay results. Direct assessment of olfaction in *Scn2a^E/+^* mice revealed intact discrimination for both social and non-social odors, but lower dishabituation to same-sex social odors compared to WT. This apparent contradiction with the hyper-social behavior observed in the three-chamber assay, which evaluates interactions with a target mouse rather than an odor- soaked cotton-swab, suggests that altered social behavior in *Scn2a^K1422E^* mice occurs downstream of olfactory perception.

Consistent with mixed effects of K1422E on channel function, *Scn2a^K1422E^* mice present with phenotypes that overlap *Scn2a* haploinsufficiency, as well as unique epilepsy-related phenotypes that are not observed in loss-of-function models. This combination of phenotypes within a single model underscores how *Scn2a^K1422E^* mice can serve as a useful platform to investigate phenotypic complexity of *SCN2A*-associated disorders.

## Acknowledgements

The genetically engineered mice were generated with the assistance of Lynn Doglio in the Northwestern University Transgenic and Targeted Mutagenesis Laboratory. This work was supported by grants NIH R21 OD025330 (JAK), NIH U54 NS108874 (ALG and JAK), SFARI 629287 and 513133 (KJB); NIH R01 MH125978 (KJB); NSF 1650113 (AML); NIH F32 MH125536 (ADN); NIH KL2TR001424 (SNM); Epilepsy Foundation of Greater Chicago Rovner Fellowship (SNM); FamileSCN2A Action Potential Grant (SNM).

